# Cryo-EM structures of SAMD9L reveal the arrangement and coordination of multi-domains

**DOI:** 10.64898/2026.02.01.703102

**Authors:** Yusuke Sekiguchi, Sushree S Sahoo, Michael L Oldham, Amanda Nourse, Marcin W Wlodarski, Ravi C Kalathur

## Abstract

Human sterile alpha motif domain-containing 9 (hSAMD9L) is a large (∼185 kDa) multi-domain interferon-stimulated antiviral effector with strong translation-inhibitory activity. Inherited heterozygous gain-of-function (GoF) mutations in SAMD9L directly associated with severe bone marrow failure syndromes. Using single-particle cryo-electron microscopy (cryo-EM), we determined the first structures of both full-length wild-type hSAMD9L and an N-terminal-truncated mutant at resolutions ranging from 2.8 to 3.7 Å. Both proteins exist in monomeric and dimeric states, providing clear evidence that the sterile alpha motif (SAM) and AlbA domains are not essential for dimerization. Our cryo-EM analysis reveals a tightly packed, closed architecture defined by interlocking multi-domains. We precisely mapped the extensive dimer interface mediated by Sir2-like and an oligonucleotide/oligosaccharide-binding (OB) domains. Biochemical analyses show that hSAMD9L binds double-stranded DNA in vitro but has no detectable NTP hydrolysis activity under our assay conditions. Accordingly, in our cryo-EM map we observed a clear density for a non-hydrolyzed NTP in the pocket, suggesting nucleotide binding without turnover like in STAND (Signal Transduction ATPases with Numerous Domains) proteins. Together, these results provide a structural framework for hSAMD9L and also providing key insights into its organization, domain packing and dimerization and offer a basis for understanding how GoF variants may alter hSAMD9L regulation thus impacting cell proliferation.

## Introduction

Sterile alpha motif domain-containing 9 (SAMD9) and SAMD9-like (SAMD9L) are interferon-inducible cytoplasmic proteins that play critical roles in antiviral immunity and negative regulation of cell proliferation (1–3). The genes encoding SAMD9 and SAMD9L arose from a tandem duplication on human chromosome 7. Most placental mammals have both paralogs (SAMD9/9L) with their coded proteins sharing approximately 60% amino acid sequence identity (1–3).

SAMD9/9L expression are transcriptionally activated by type I interferon signaling and both act as key restriction factors against DNA viruses, such as vaccinia and monkeypox (3–5), and RNA viruses, such as HIV-1(6) and COVID-19 (7). Their antiviral effects are found in multiple mechanisms, including binding to double-stranded DNA and RNA (8), association with cytoplasmic granules (4, 9–11), and cleavage of specific tRNAs, notably tRNA^Phe^, via the endoribonuclease activity of the AlbA domain (12, 13). These functions contribute to translational repression and antiviral responses (6, 14); consequently, SAMD9/9L are antagonized by several virus-encoded host-range proteins (3, 15, 16). Prokaryotic structural analogs of SAMD9/9L have been recently identified in the Avs anti-phage system; these bacterial analogs harbor predicted nuclease active sites essential for cell death, suggesting convergent evolution of immune defenses across kingdoms (17).

Clinically, gain-of-function (GoF) mutations in SAMD9/9L have been associated with myelodysplastic syndrome, bone marrow failure, ataxia-pancytopenia syndrome, and SAMD9/9L-associated autoinflammatory disease (18–24). In some patients, negative selection against GoF mutations results in somatic loss of the mutant allele from chromosomal deletion (e.g., monosomy 7) (21, 25). Many GoF mutations result in growth arrest and defective hematopoiesis, yet the molecular basis by which these mutations alter SAMD9/9L function remains unclear.

SAMD9/9L are large multidomain proteins that contain in order, a sterile alpha motif (SAM) domain, a flexible linker region (FL), an AlbA domain, a variant Sir2-like domain, a STAND-like P-loop NTPase, a tandem tetratricopeptide repeat (TPR) domain, and an oligonucleotide/oligosaccharide-binding (OB) fold domain (26–28). Their complex architecture suggests that SAMD9/9L may be capable of responding to intracellular stress or infection through structural rearrangements and functional modulations (3, 24). Several studies have demonstrated infection-dependent activation, derepression, or relocalization of SAMD9/9L, supporting that these proteins might exist in a closed or inaccessible state that becomes activated upon viral infection (3, 8, 11, 12, 27). Upon infection, SAMD9/9L form antiviral granules in the cytoplasm, which repress the translation of viral mRNA; such granules are absent without infection or when counteracted by viral antagonists (4, 10, 11). Some germline GoF mutations in SAMD9L can globally suppress protein synthesis via inhibition of translation elongation (22).

Despite roles for SAMD9/9L in antiviral immunity and negative regulation of cell proliferation, their overall molecular architectures have remained unknown. While the SAM and AlbA domains have been characterized in isolation (27, 28), how the structural relationships between all domains in the full-length protein leads to regulation have not been resolved.

In this study, we present the structural analysis of full-length wild-type human SAMD9L (hSAMD9L) and a truncated mutant lacking the N-terminal SAM domain and flexible linker region (**Δ**SAM-FL). Using cryo-EM, biophysical characterization, and functional assays, we demonstrate that hSAMD9L exists as both monomeric and dimeric species in solution and adopts a compact, domain-packed conformation. Our findings provide a framework for understanding how domain organization controls hSAMD9L function and for dissecting the structural consequences of disease-associated mutations.

## Results

### Expression and purification of hSAMD9L

For our initial expression screening, we designed four constructs (Figure 1A): hSAMD9L comprising residues 2–1584 of wild-type hSAMD9L; ΔSAM hSAMD9L (residues 80–1584) representing a truncation of the N-terminal SAM domain; ΔSAM-FL hSAMD9L (residues 155–1584) lacking both the N-terminal SAM domain and the flexible linker region; and ΔOB hSAMD9L (residues 2–1178) representing a C-terminal OB domain truncation.

**Figure 1.**
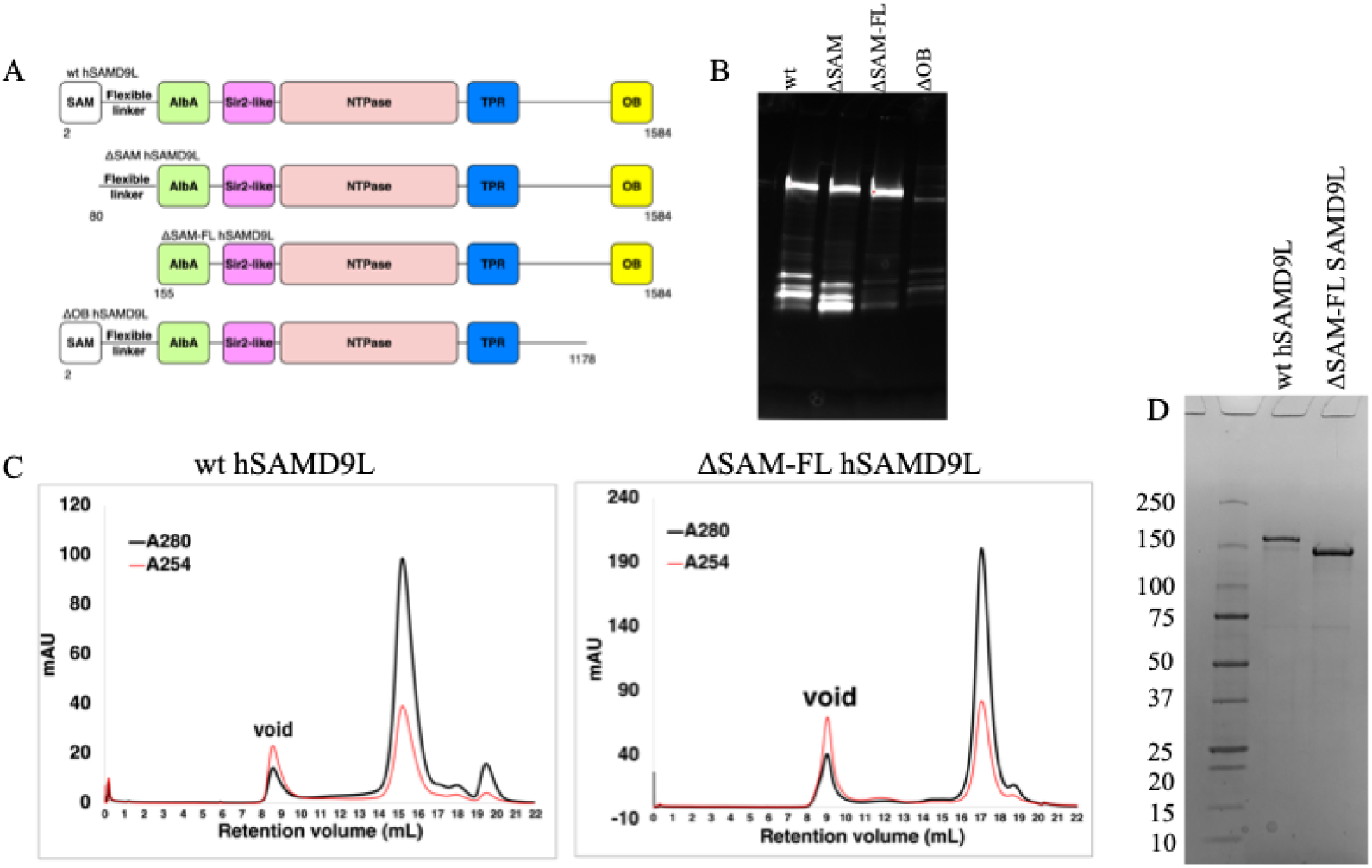
Constructs and expression screening of GFP-tagged hSAMD9L variants. **A**: Schematic representation of four constructs designed for expression screening. From top to bottom: wt (residues 2–1584), **Δ**SAM (residues 80–1584), ΔSAM-FL (residues 155–1584), and ΔOB (residues 2–1178). **B**: In-gel GFP fluorescence image showing the expression of each construct. Lanes from left to right: wt, ΔSAM, ΔSAM-FL, and **Δ**OB. The upper bands correspond to full-length GFP–hSAMD9L fusion proteins, while the lower bands represent degraded products. **C:** Size-exclusion chromatography (SEC) profiles of wt hSAMD9L and ΔSAM-FL hSAMD9L using a Superose 6 10/300 column. Elution was monitored at two wavelengths: A280 (black line) and A254 (red line). **D**: SDS-PAGE analysis of the final purified samples. Lanes from left to right: wt hSAMD9L (184.9 kDa) and ΔSAM-FL hSAMD9L (167.3 kDa).

All constructs were expressed as N-terminal GFP fusion proteins in *Expi293 GnTI⁻*cells, and expression levels were assessed by GFP fluorescence on SDS-PAGE (Figure 1B). Both hSAMD9L and ΔSAM hSAMD9L exhibited bands at the expected molecular weights, along with some degradation products. In contrast, ΔSAM-FL hSAMD9L showed no detectable degradation, and ΔOB hSAMD9L displayed suboptimal expression levels.

Based on these results, we selected hSAMD9L and ΔSAM-FL hSAMD9L for purification and analysis. To reduce nucleic acid contamination, benzonase was added before cell lysis. Both exhibited favorable monodispersity and homogeneity as assessed by size-exclusion chromatography and SDS-PAGE (Figure 1C and 1D).

### Purified hSAMD9L binds to dsDNA

Full-length SAMD9 has previously been reported to bind double-stranded DNA (dsDNA), thereby inducing its oligomerization (8). Also, it was reported that the truncated SAMD9 without both AlbA and OB domains could still bind dsDNA (8). To determine whether the purified hSAMD9L proteins would also exhibit behavior similar to that of SAMD9, we performed electrophoretic mobility shift assays (EMSA) and native PAGE analysis. EMSA showed that both wt full-length hSAMD9L and the ΔSAM-FL hSAMD9L can bind dsDNA (Figure 2). Native PAGE further revealed that dsDNA binding promoted oligomerization or aggregation of hSAMD9L (Suppl figure 1).

**Figure 2.**
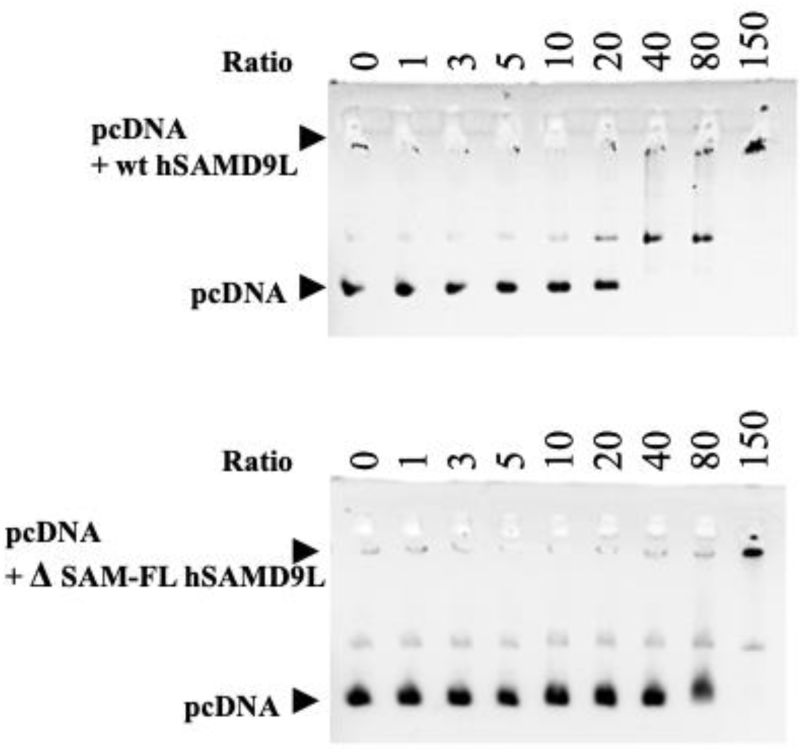
Electrophoretic mobility shift assay (EMSA) of circular plasmid dsDNA with increasing amounts of hSAMD9L. EMSA was performed using circular plasmid DNA (pcDNA) incubated with increasing concentrations of hSAMD9L (molar ratio from 0 to 150). The lower arrow indicates free pcDNA, while the upper arrow indicates the DNA–protein complex. **Top**: wt hSAMD9L. **Bottom**: ΔSAM-FL hSAMD9L.

### wt hSAMD9L adopts monomeric and dimeric states

In cryo-electron microscopy (cryo-EM), we observed two distinct particle populations in our 2D classifications and 3D reconstructions representing both the monomeric and dimeric forms (Suppl figures 2 and 19). To determine whether the observed dimer was an artifact of cryo-EM, we conducted additional biophysical analyses. The size-exclusion chromatography (SEC) profile displayed a single, sharp Gaussian peak, making it impossible to distinguish between the monomeric and dimeric forms using SEC alone. Therefore, we turned to mass photometry for further evaluation.

The mass photometry analysis indicated that at a concentration of 100 nM, both wild-type hSAMD9L and the ΔSAM-FL variant primarily exist as monomers, with a smaller population of dimers (Suppl figure 3). Since mass photometry experiments are typically conducted at very dilute concentrations, this confirmed the presence of dimers in solution. Furthermore, the presence of a dimeric form for ΔSAM-FL hSAMD9L suggests that neither the SAM domain nor the flexible linker is necessary for dimer formation.

To evaluate the concentration-dependent oligomer formation, we performed sedimentation velocity analytical ultracentrifugation (SV-AUC) at higher concentrations (500 nM for hSAMD9L; 1.0 µM for ΔSAM-FL). The proteins partitioned between monomer and dimer, without evidence of higher-order oligomers (Suppl figure 4, Table 1 and 2). Together, these data suggest that hSAMD9L predominantly exists as monomer and dimer over the tested concentration range (0.1–1.0 µM) and did not observe multimers beyond dimers.

**Table 1.** 1D Sedimentation velocity – analytical ultracentrifugation (SV-AUC): Summary of results of the sedimentation velocity *c(s)* analysis of wt hSAMD9L and ΔSAM-FL hSAMD9L.

| *Protein construct* | *s<sub>20</sub> (Svedberg)* | *M<sub>w</sub> (Da)* | *(f/f<sub>0</sub>)<sub>w</sub>* |
| --- | --- | --- | --- |
| wt hSAMD9L | 6.67 (30%) | 185,057 | 1.50 |
|  | 9.12 (9%) | 296,028 | 1.50 |
|  | 2.86 (37%) | 52,116 | 1.50 |
|  | 4.95 (20%) | 118,386 | 1.50 |
| ΔSAM-FL hSAMD9L | 6.75 (40%) | 171,916 | 1.40 |
|  | 10.01 (13%) | 310,464 | 1.40 |
|  | 2.54 (18%) | 39,737 | 1.40 |
|  | 4.67 (12%) | 98,875 | 1.40 |

Weight-average sedimentation coefficient calculated from $c(s)$ distribution at 20 °C with percentage of total protein indicated. Molar mass values (MW) taken from the $c(s)$ -transformed $c(M)$ distribution. Best-fit weight-average frictional ratio values $(f/f_0)_w$ taken from the $c(s)$ distribution.

**Table 2.**
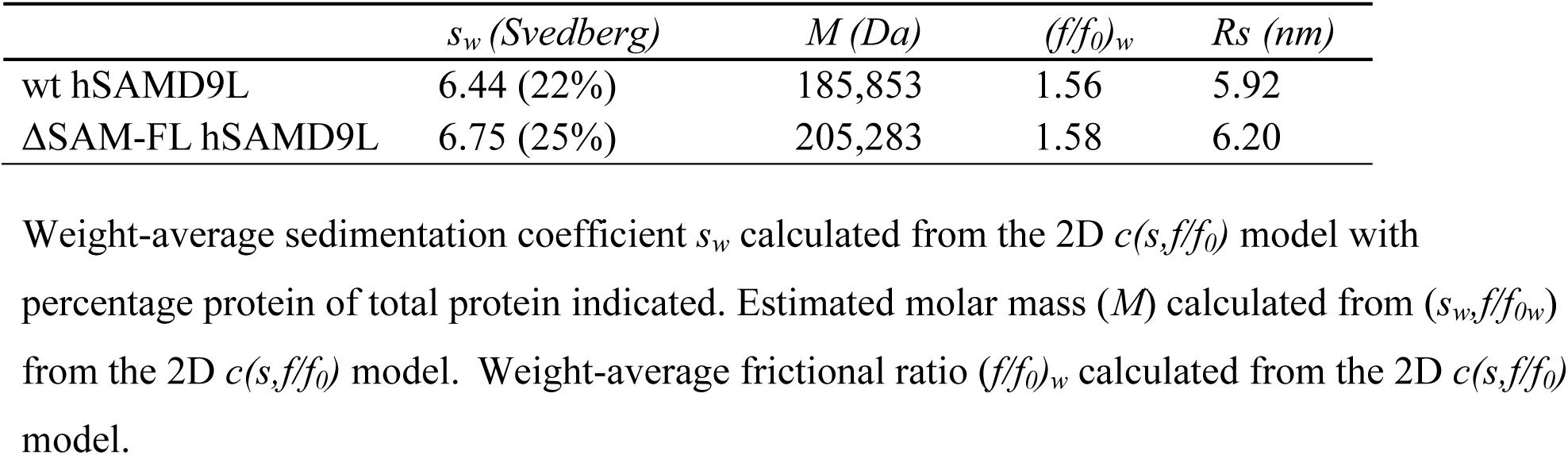
Best-fit values and estimates of the two-dimensional *c(s,f/f_0_*) analyses of wt hSAMD9L and ΔSAM-FL hSAMD9L.

### wt hSAMD9L monomer adopts a compact conformation

Our cryo-EM reconstructions for both monomeric and dimeric forms of hSAMD9L yielded resolutions of 2.84 Å and 2.89 Å, respectively (Suppl figures 5, 6 and 19). The concentration of hSAMD9L used for cryo-EM was 6.5 µM and the number of particles observed was greater for the monomer than for the dimer (Table 3), in agreement with the mass photometry and SV-AUC data.

**Table 3.**
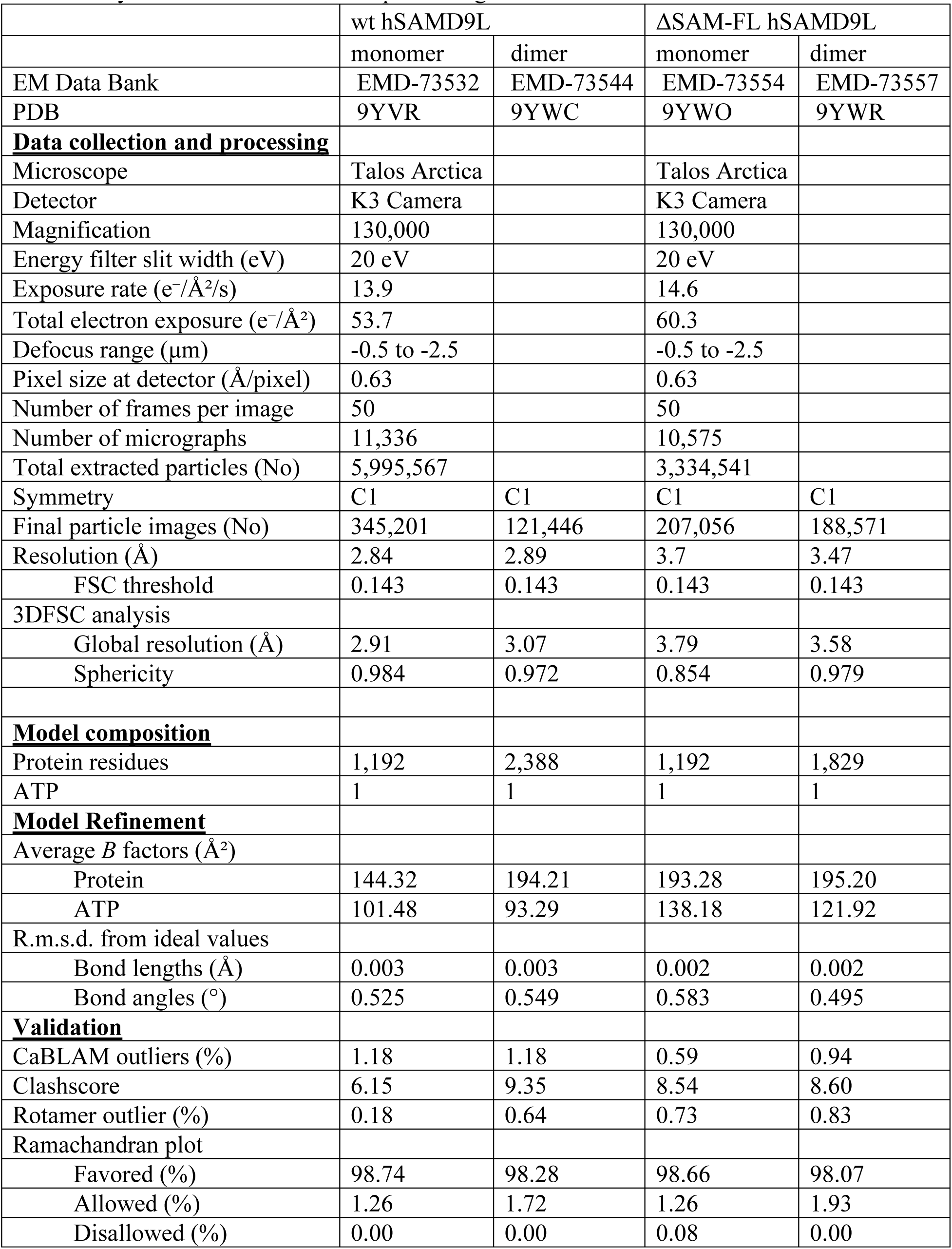
cryo-EM data collection and processing statistics.

For the monomeric form, residues 393–1584 were assigned to the cryo-EM map, corresponding to the Sir2-like, NTPase, TPR, and OB domains (Figure 3A). We identified a newly structured region between the TPR and OB domains, composed of multiple α-helices, which we will refer to as the Helical domain. In contrast, the N-terminal region (residues 2–392), which includes the SAM domain, flexible linker region, and AlbA domain, is disordered with no observable densities.

**Figure 3.**
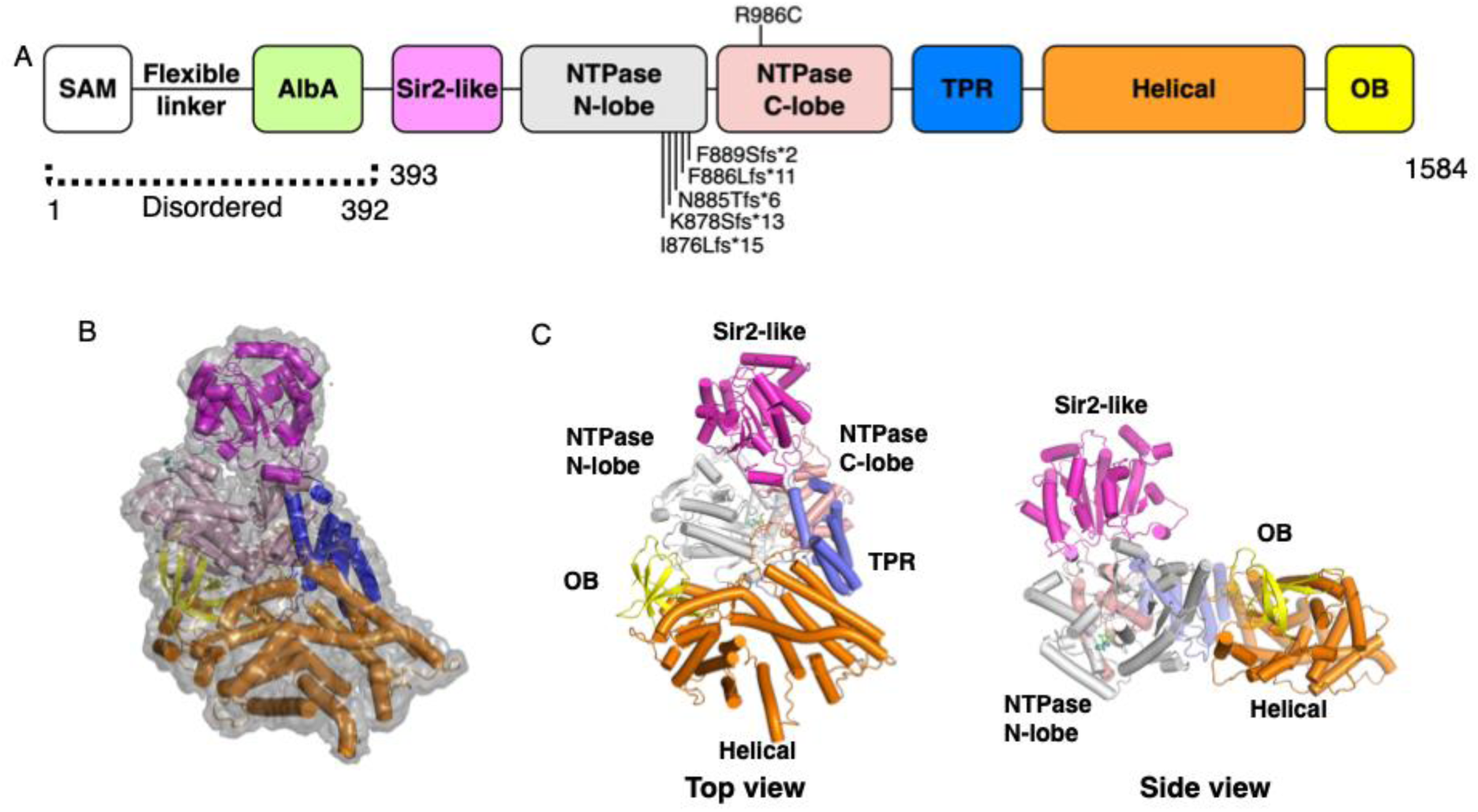
cryo-EM structure of wt hSAMD9L A: Domain organization of hSAMD9L. Schematic diagram of the full-length structure showing annotated domains. The region from the SAM to the AlbA domain is not resolved in the map. The NTPase domain is divided into N-lobe and C-lobe for clarity. Disease-associated missense mutation and frameshift mutations are mapped to the NTPase domain. The region between the TPR and OB domains is designated as the “helical domain.” **B: Cryo-EM map and structural model.** cryo-EM maps of the wt hSAMD9L monomer is shown, with the fitted atomic models represented as ribbon diagrams. **C: Ribbon representation of the wt hSAMD9L structure.** The structure of wild-type hSAMD9L is shown as a ribbon model. Each domain is color-coded according to the scheme used in the construct schematic.

The ordered domains pack tightly to form a compact structure. Its architecture resembles a foot, with the Sir2-like domain as the ankle, the NTPase domain as the heel, the TPR domain forming the arch, and the Helical and OB domains representing the toes. The Helical and OB domains interact with the N-lobe of the NTPase domain, contributing to the compact conformation. The relatively lower local resolution of the Helical and OB domains may indicate intrinsic structural flexibility (Suppl figure 5).

The NTPase domain consists of two lobes: the N-lobe, which contains a P-loop NTPase domain (26), and the C-lobe (Suppl figure 7). The P-loop NTPase domain harbors conserved motifs, including Walker A, Walker B, and the arginine finger (Suppl figure 8). Notably, the Walker A motif contains His741 instead of the conserved lysine residue typically required for ATP hydrolysis, which may compensate for the catalytic function (Figure 4) (29–32). A clear density was observed in cryo-EM map in the ATP-binding pocket, we could confidently build a purine nucleotide (Suppl figure 7), but resolution is not good enough to differentiate if it is an adenine or guanine. What is unambiguous is the strong density for the γ-phosphate, confirming a unhydrolyzed (intact) NTP, we provisionally modeled an ATP into the density that presumably originated from host cells given that no nucleotide was added during purification. To assess whether the purified sample has intrinsic NTPase activity, we performed ATP/GTPase assays using wt hSAMD9L but detected no ATPase or GTPase activity under our experimental conditions (Suppl figure 9).

**Figure 4.**
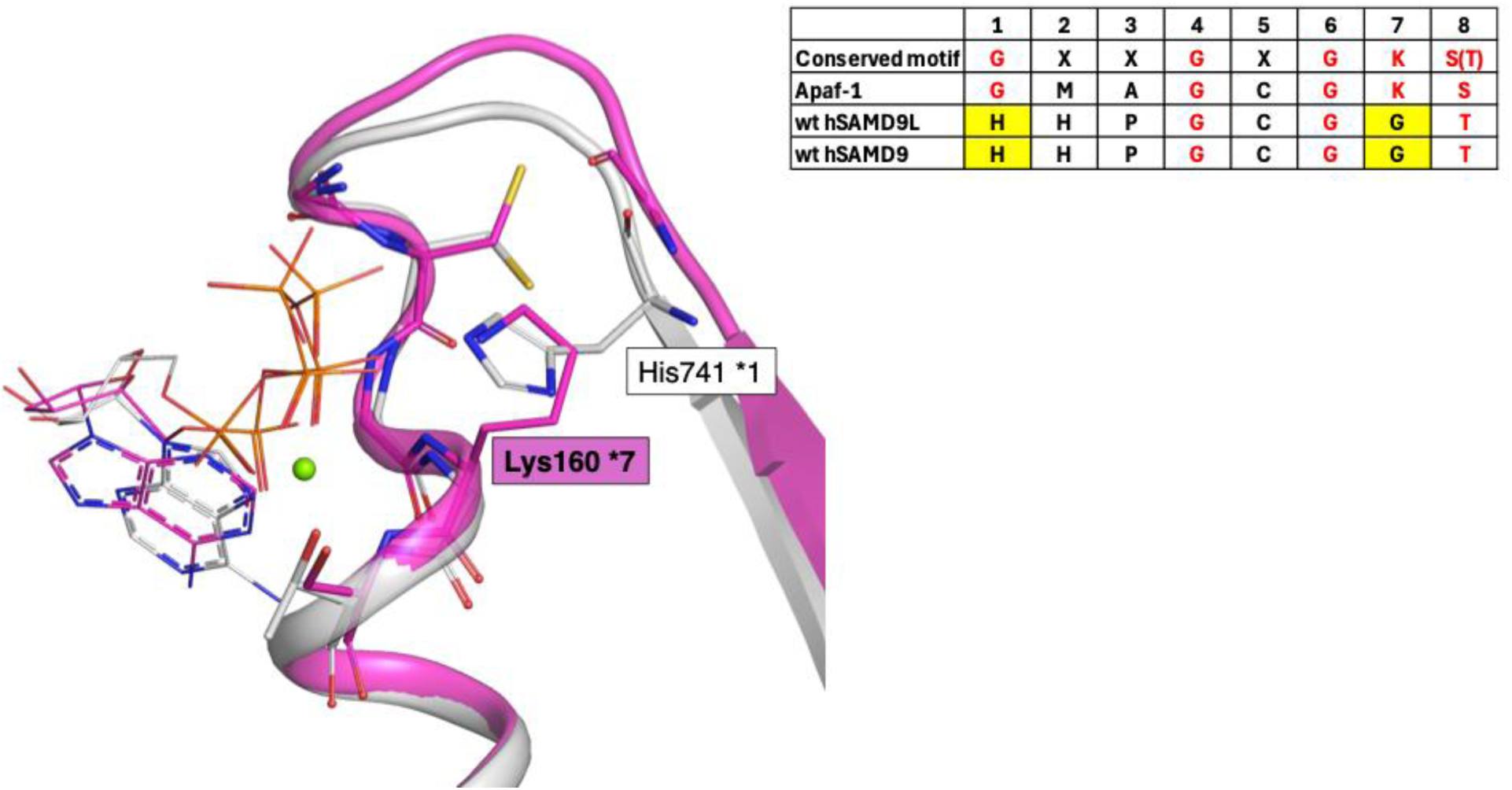
Structural comparison of the ATPase motifs between wt hSAMD9L and Apaf-1. In hSAMD9L, the nitrogen atom of a histidine residue located on the opposite side replaces the conserved lysine side chain nitrogen typically found in P-loop ATPases. **White**: wt hSAMD9L **Purple**: Apaf-1 (PDB ID: 3JBT)

### wt hSAMD9L dimer is in a closed (interlocked) conformation

The dimeric wt hSAMD9L structure shows some differences from the monomeric form. Similar to the monomer, both protomers within the dimer also exhibited disordered SAM domains, flexible loops, and AlbA domains, (Figure 5 and Suppl figure 10 A, B). One protomer (protomer A) adopted a conformation nearly identical to that of the monomeric form and shows clear cryo-EM density for a nucleotide in the NTP-binding site (Suppl figure 11). In contrast, the second protomer (protomer B) displayed a substantially altered conformation in which the Sir2-like domain was rotated away from the NTPase domain (Figure 5 and Suppl figure 10 B, C). Additionally, the Helical domain was split at Leu1310 into two segments (Helical-A and Helical-B), with Helical-B and the OB domain further away from the NTPase domain than in the monomer or protomer A. Because of the limited local resolution, we cannot make an assessment about the presence of nucleotide in the NTPase domain of protomer B.

**Figure 5.**
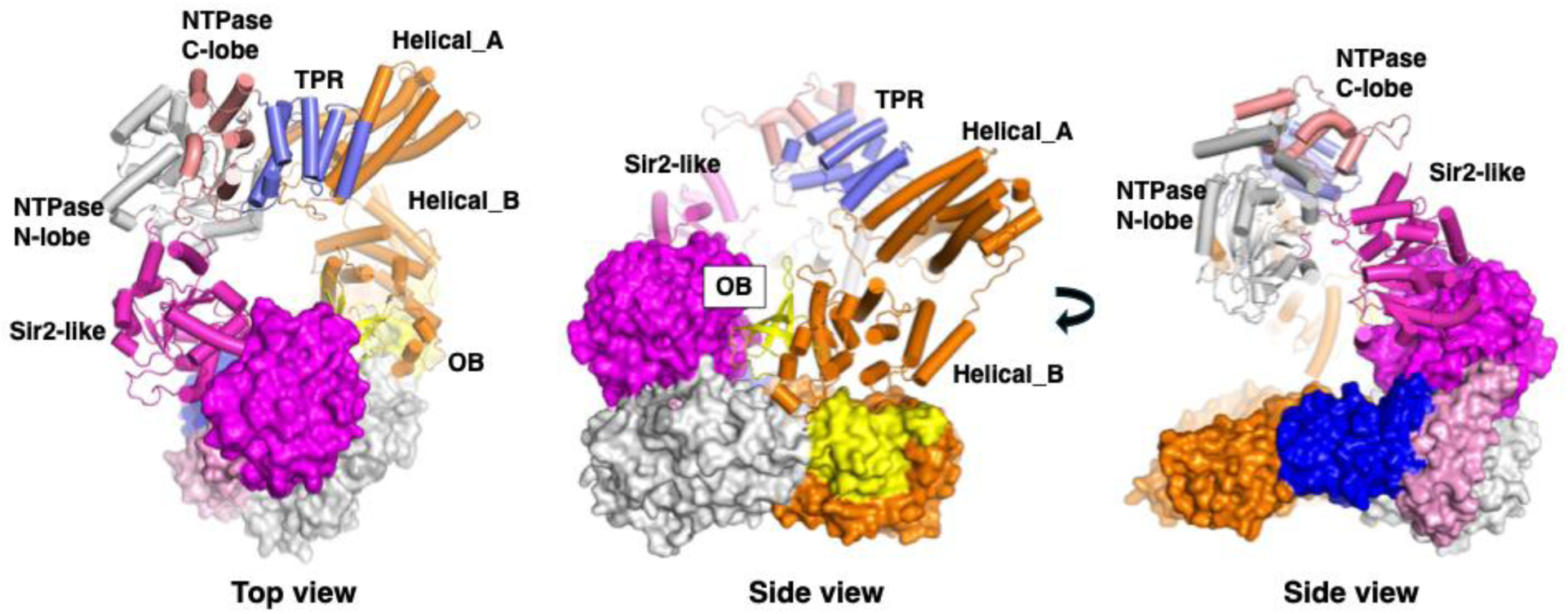
Dimeric structure of wt hSAMD9L. The protomer A is shown in surface representation, while the protomer B is shown as a ribbon model. The protomer B interacts with the protomer A through its Sir2-like and OB domains.

There are two major dimer interfaces. The first interface involves the dimerization of the Sir2-like domains, mediated by hydrogen bonds and salt bridges between multiple residue side chains (Suppl figure 12 and Table 4). According to PISA server analysis (https://www.ebi.ac.uk/pdbe/pisa/) (33), this interface covers a buried surface area of 1017.6 Å². In the cryo-EM map, protomer B generally had lower resolution than protomer A, but the Sir2-like domains of both protomers show comparable resolution, suggesting structural rigidity in this region due to the interdomain interactions (Suppl figure 6). Sir2-like domains in other proteins are also known to mediate oligomerization (34–36). The second dimer interface is formed with extensive contacts between the Sir2-like domain of protomer A and the OB domain of protomer B (Suppl figure 13 and Table 5). The surface area of this interface was calculated to be 1295.2 Å² according to PISA analysis.

**Table 4.** Sir2-like–Sir2-like interaction interface in the dimeric structure of wt hSAMD9L.

**/HYDROGENBONDS**
| **/STRUCTURE/DISTANCE** | **/STRUCTURE/STRUCTURE1** | **/STRUCTURE/STRUCTURE2** |
| --- | --- | --- |
| 2.4 | B:GLN 569[ NE2] | A:GLN 518[ OE1] |
| 3.6 | B:ARG 597[ NH2] | A:GLN 518[ OE1] |
| 2.9 | B:LYS 476[ NZ ] | A:ASN 536[ OD1] |
| 3.4 | B:GLN 569[ NE2] | A:GLU 562[ OE1] |
| 3.2 | B:ARG 597[ NH2] | A:GLU 562[ OE1] |
| 2.6 | B:ARG 519[ NE ] | A:TYR 568[ OH ] |
| 2.9 | B:GLU 562[ OE2] | A:GLN 569[ NE2] |

**Table 5.** Sir2-like–OB interaction interface in the dimeric structure of wt hSAMD9L.

| **/STRUCTURE/DISTANCE** | **/STRUCTURE/STRUCTURE1** | **/STRUCTURE/STRUCTURE2** |
| --- | --- | --- |
| 3.6 | B:HIS1508[ NE2] | A:ALA 789[ O ] |
| 3.5 | B:LYS1532[ NZ ] | A:GLU 779[ OE2] |
| 3.8 | B:TYR1552[ OH ] | A:GLU 824[ O ] |
| 2.7 | B:SER1553[ OG ] | A:GLU 779[ OE2] |
| 2.7 | B:ARG1557[ NH1] | A:ASN 466[ OD1] |
| 3.6 | B:ARG1557[ NH1] | A:GLN 492[ OE1] |
| 3.5 | B:ARG1557[ NH2] | A:ALA 465[ O ] |
| 2.9 | B:SER1558[ OG ] | A:GLU 776[ OE2] |
| 2.9 | B:ILE1562[ N ] | A:LEU 467[ O ] |
| 2.4 | B:GLU1530[ OE1] | A:GLN 472[ NE2] |
| 3.3 | B:GLY1559[ O ] | A:ASN 466[ ND2] |
| 3.0 | B:GLY1559[ O ] | A:HIS 468[ NE2] |
| 3.6 | B:ARG1560[ O ] | A:GLN 472[ NE2] |
| 3.0 | B:ASN1561[ OD1] | A:LEU 467[ N ] |
| 2.4 | B:ILE1562[ O ] | A:ARG 501[ NH1] |
| 3.9 | B:GLU1563[ OE1] | A:ARG 501[ NH2] |
| 3.6 | B:GLU1563[ OE2] | A:LYS 460[ NZ ] |

### Structure of ΔSAM-FL (truncated) hSAMD9L

We also determined the structure of the **Δ**SAM-FL construct, which also exists in monomeric and dimeric forms in solution (Suppl figures 3 and 4). Cryo-EM data were processed for both monomeric and dimeric forms, yielding resolutions of 3.70 Å and 3.47 Å, respectively (Suppl figures 14, 15 and 20 Table 3). As observed for full-length hSAMD9L, the AlbA domain was disordered in both monomeric and dimeric forms. In the dimer form, residues beyond position 1308 in protomer A and residues 1096–1337 in protomer B were also disordered (Suppl figure 16).

The structure of monomeric **Δ**SAM-FL hSAMD9L shows architecture that is essentially identical to that of full-length hSAMD9L. In the dimer structure, protomers A are also very similar. In contrast, protomer B showed differences in the positioning of the NTPase domain, TPR domain, and the N-terminal portion of the Helical domain. In the **Δ**SAM-FL dimer, these domains were shifted closer to the protomer A (Suppl figure 17) than in hSAMD9L. This observation, along with the lower local resolution of those regions in both the wt and **Δ**SAM-FL structures, suggests that these domains possess some conformational flexibility. However, the conserved arrangement of the Sir2-like domain and the C-terminal portion of the Helical and OB domains between the two protomers supports the idea that these regions contribute critically to dimer interface stabilization.

Importantly, the structures of **Δ**SAM-FL indicate that the SAM domain and flexible loop region were not required for the folding, stability and architecture of either the monomeric or dimeric forms. These domains may instead play a role in higher-order oligomerization of SAMD9L, possibly in response to viral infection or other stimuli.

## Discussion

Our structural, mass photometry, and AUC analyses of both full-length and **Δ**SAM-FL confirmed that the SEC-purified protein sample contained a mixture of monomers and dimers, but no other multimers; the dimer is real and not an experimental artifact. Our study is the first cryo-EM structure to provide atomistic and architectural details of the monomeric and dimeric forms, with both monomers adopting a compact architecture and the dimers adopting a compact interlocked architecture. Although the isolated SAM domain of SAMD9 can form a polymer (56), in our structure, this domain is disordered. Notably, in both monomeric and dimeric structures of full-length hSAMD9L, surprisingly, the AlbA domain is also disordered, even though it is the only domain with demonstrated catalytic activity (tRNA cleavage) and is not connected by a flexible linker, unlike the Sam domain.

Based on these observations, we propose that the compact conformation visualized here likely represents a locked or inaccessible conformation of hSAMD9L. Such a locked state may represent a regulatory mechanism by which hSAMD9L activity is controlled. Given that overactivation of hSAMD9L in normal cells can lead to detrimental effects, such as inhibition of cell proliferation (2, 25), this inaccessible conformation may serve as a safeguard to prevent excessive hSAMD9L activity under non-stimulated conditions. Because hSAMD9L is constitutively expressed even in the absence of viral infection (1), unregulated activation under basal conditions could undesirably suppress cell proliferation. Therefore, maintaining this inaccessible conformation is likely critical for preventing unintended cellular consequences.

Our observations also indicate that protomer B in the dimer undergoes significant conformational rearrangement relative to the protomer A and that dimer stabilization is mediated through inter-protomer interactions between the Sir2-like and OB domains. This conformational shift may represent an intermediate state in the transition of hSAMD9L from a closed (inaccessible), domain-packed conformation to a more open, domain-relaxed form. Despite this domain movement, the dimer maintains an overall compact architecture, suggesting that domain release (or opening) occurs in a coordinated yet constrained manner.

Although the precise relationship between oligomerization and hSAMD9L’s functional activity remains unclear, our structural analysis revealed that the AlbA domain, the only currently known catalytic domain of SAMD9 (12), is disordered and likely exposed to solvent. These findings suggest that hSAMD9L may exhibit catalytic activity even in its closed conformation or that binding to tRNA induces a transition from the closed to the open state.

The recurrent GoF missense variant p.Arg986Cys (R986C) is associated with ataxia-pancytopenia syndrome and myelodysplastic syndrome; functional studies indicate enhanced antiproliferative/translation-repressive activity, consistent with a pathogenic GoF mechanism. (5, 9, 24, 37). Our structures show that Arg986 is in the C-lobe of the P-loop NTPase domain, with its side chain oriented toward the N-lobe and forming a hydrogen bond with the backbone carbon of Gly690. Substitution of Arg986 with cysteine would disrupt this interaction and weaken the interaction between the N- and C-lobes (Suppl figure 18). Notably, most reported GoF mutations predominantly cluster around the NTPase domain, particularly frameshift mutations are found in downstream of the N-lobe (Figure 3A) (23, 24). These mutations result in loss of the C-lobe, which is responsible for shielding the NTP-binding site. These observations suggest that the P-loop NTPase domain plays a crucial role in maintaining the closed conformation and structural stability of hSAMD9L, along with the tight assembly of the TPR, Helical, and OB domains. Conformational changes induced by domain dissociation may trigger activation and oligomerization. Since in our AUC experiments at high concentrations, we predominantly saw monomers and have not observed multimers beyond dimers, this suggest that there is considerable energy barrier for the transition to multimeric or open form and so that may be reason why there are lot of GoF mutations mapped in this region.

A recent study uncovered a link between ancient immune defense mechanisms in prokaryotes and evolutionary adaptations to lentiviral challenges in primates (38). Members of the signal transduction ATPases with numerous domains (STAND) superfamily are typically multidomain proteins composed of a sensor, an NTPase, and an effector domain. Pathogen recognition by the sensor domain triggers oligomerization, which in turn induces effector activity (39). Sequence and structural analyses indicate that SAMD9L has a domain architecture consistent with this paradigm (26). Although hSAMD9L has conserved Walker A/B motifs and a clear unambiguous density in cryo-EM map corresponding to an intact NTP, we did not detect nucleotide hydrolysis activity in our preparations. In STAND members, nucleotide binding often acts as the switch for large-scale conformational changes and higher-order assembly, whereas the requirement for hydrolysis is system-dependent. For example, in the case of the STAND NTPase that assembles the apoptosome, Apaf-1, binding of dATP/ATP promotes oligomerization; yet, hydrolysis is not strictly required (40). Thus, it is possible that for hSAMD9L, nucleotide binding is the primary determinant of conformational change and assembly of higher-order oligomeric states.

In prokaryotic Avs/STAND systems and the SIR2-containing protein defense pathway, pattern recognition of viral factors initiates assembly/oligomerization, followed by allosteric induction of effector activities such as Sir2 (NADase) (34–36, 41). In addition, Sir2 domains can also contribute structurally as assembly scaffolds, as reported for bacterial SIR2-containing proteins (34–36), Avs5 (42), and Apaf-1 (40). In the case of hSAMD9L Sir2-like domain, no classical sirtuin (deacetylase) activity has been reported, and the canonical Cys₄-type Zn²⁺ binding domain found in many sirtuins is absent (43). These observations suggest that hSAMD9L Sir2-like domain has a primarily regulatory/scaffolding role rather than an enzymatic one, whereas the AlbA domain’s tRNA cleavage activity is the major antiviral effector output (12).

Taken together, we propose a division of labor model in which the Sir2-like, STAND-like NTPase and OB domains of hSAMD9L tune assembly states (conformational control and higher-order oligomerization), whereas the AlbA domain constitutes the principal antiviral effector. Future work will be required to define the structures of activated or higher-order oligomeric states, characterize DNA-bound assemblies, and determine how GoF and loss-of-function variants disrupt regulatory architecture.

## Methods

### Construct screening

Wild-type hSAMD9L (residues 2–1584), an N-terminally truncated mutant (residues 80–1584), a construct lacking both the N-terminal SAM domain and the flexible linker (residues 155–1584), and a C-terminally truncated mutant (residues 2–1178) were cloned into the *pEG BacMam* vector. Each construct was fused at the N-terminus with a 10×His tag, mEGFP, and an HRV 3C protease cleavage site.

*Expi293 GnTI⁻* cells were transiently transfected with each plasmid by using the *ExpiFectamine™ 293 Transfection Kit*, and cells were harvested after 2–3 days of incubation. Harvested cells were lysed by sonication, and cell debris was removed by centrifugation. The resulting supernatants were analyzed by SDS-PAGE without pre-heating and imaged based on GFP fluorescence.

### Large-scale expression and purification

The wt hSAMD9L gene (residues 2–1584) was cloned into the pcDNA3.4 vector with an N-terminal 3×FLAG tag and an HRV 3C protease cleavage site. For **Δ**SAM-FL hSAMD9L (residues 155–1584) expression, we used the same construct as used in the construct screening. *Expi293 GnTI⁻*cells were transiently transfected with each plasmid using the *ExpiFectamine™ 293 Transfection Kit* and harvested after 2–3 days of incubation. For wt hSAMD9L purification, harvested cells were resuspended in buffer containing 20 mM HEPES-NaOH (pH 7.4), 300 mM NaCl, protease inhibitor cocktail, and benzonase, then lysed by sonication. Cell debris was removed by centrifugation, and the supernatant was incubated with *anti-FLAG M2 resin* pre-equilibrated with the same buffer. The resin was gently rotated for 1 h and loaded onto an open column. After washing with 30 column volumes (CV) of buffer, the bound protein was eluted using buffer containing FLAG peptide. Eluted fractions were concentrated and subjected to size-exclusion chromatography using a Superose 6 Increase 10/300 column. Peak fractions were pooled, concentrated to 1.2 mg/mL, flash-frozen in liquid nitrogen, and stored at −80 °C.

For **Δ**SAM-FL hSAMD9L purification, cells were processed similarly. After lysis in the same buffer, the supernatant was incubated with anti-GFP nanobody resin pre-equilibrated with buffer. Following 1 h of rotation, the resin was washed with 30 CV of buffer, and HRV 3C protease was added directly to the resin for on-resin cleavage. After 1 h of incubation, the flow-through was collected, concentrated, and applied to a Superose 6 Increase 10/300 column. Peak fractions were pooled, concentrated to 0.8 mg/mL, flash-frozen in liquid nitrogen, and stored at −80 °C.

### Mass photometry

Mass photometry was performed by using a Refeyn instrument. A microwell was prefilled with buffer containing 20 mM HEPES-NaOH (pH 7.4) and 300 mM NaCl, and purified wt hSAMD9L or **Δ**SAM-FL hSAMD9L was added directly to the well to achieve a final protein concentration of 100 nM. Data were acquired as 6,000-frame videos (60 s) and analyzed using DiscoverMP software-v2023 R.1.2 (Refeyn).

### Sedimentation velocity – analytical ultracentrifugation

Sedimentation velocity experiments were conducted in a ProteomeLab XL-I analytical ultracentrifuge (Beckman Coulter, Indianapolis, IN) with an AnTi-50 eight-hole rotor following standard protocols unless mentioned otherwise (44). The sample in 50 mM HEPES-NaOH, pH 7.4, 300 mM NaCl buffer was loaded into cell assemblies with double sector 12 mm centerpieces and sapphire windows. The cell assembly, containing identical sample and reference buffer volumes of 200 µL, was placed in a rotor. After temperature equilibration at 20 °C, the rotor was accelerated to 42,000 rpm, and Rayleigh interference optical data were collected. The velocity data were analyzed via the sedimentation coefficient distribution model *c(s)* in SEDFIT (http://sedfitsedphat.nibib.nih.gov/software) (45). The signal-average frictional ratio and meniscus position were refined via non-linear regression, and maximum entropy regularization was applied at a confidence level of P-0.68. Buffer density and viscosity were measured at 20 °C by using a densitometer (Anton Paar Inc., model DMA 5000 M) and a micro-viscometer (Anton Paar Inc., model AMVn), respectively. The partial specific volumes and the molecular masses of the proteins were calculated based on their amino acid compositions in SEDFIT (http://sedfitsedphat.nibib.nih.gov/software).

The sedimentation velocity data were also analyzed with the two-dimensional size-shape distribution, *c*(*s,f/f_0_*) model (with one dimension shown by the *s-*distribution and the other shown by the *f/f_0_*-distribution). The equidistant *f/f_0_*-grid of 0.25 steps varied from 0.5 to 2.5, and the linear *s*-grid of 100 s-values from 2 to 20 S. Tikhonov-Phillips regularization was at one standard deviation. The velocity data were then transformed to *c(s,f/f_0_), c(s,M) and c(s,R)* distributions (with *M* = molar mass, *f/f_0_* = frictional ratio, R = Stokes radius, and s = sedimentation coefficient), and plotted as contour plots. The color temperature of the contour lines indicates the population of species (44, 46). All plots were created in GUSSI (http://www.utsouthwestern.edu/labs/mbr/software/). (47)

### EMSA assay

Ten ng of pcDNA3.4 plasmid was mixed with either wt hSAMD9L or **Δ**SAM-FL hSAMD9L at molar ratios ranging from 1:1 to 1:150 in buffer containing 300 mM NaCl, 5 mM MgCl₂, 1 mM DTT, and 10% glycerol. The mixtures were incubated for 1 h at 30 °C.

Each sample was then loaded onto a 0.8% agarose gel and electrophoresed in 0.5X TAE buffer at 50 V for 60 min. After electrophoresis, the gel was stained with GelRed for 30 min and imaged by using ChemiDoc imaging system (BioRad).

### Native-PAGE

Circular pcDNA3.4 plasmid DNA (0.17 µM) was mixed with 1 µM wt hSAMD9L or **Δ**SAM-FL hSAMD9L in buffer containing 50 mM HEPES–NaOH (pH 7.4), 300 mM NaCl, and 5 mM MgCl₂. The mixtures were incubated for 1 h at 30 °C. Samples were combined with Quick Fluorescent Loading Buffer (MP Biomedicals) according to the manufacturer’s instructions, loaded onto native polyacrylamide gels, and electrophoresed at 20 mA for 1 h at 4 °C. Gels were imaged under UV illumination.

### ATP/GTPase activity assay

ATP/GTPase activity was measured by using the Malachite Green Phosphate Assay Kit (Sigma-Aldrich, MAK113). Purified wt hSAMD9L was tested at concentrations ranging from 0.03 μM to 33.75 μM. Samples were mixed with reaction buffer containing 4 mM ATP or GTP and incubated at room temperature for 1 h. As a positive control, apyrase was used at concentrations ranging from 0.25 mU to 250 mU. Reactions were terminated by adding detection reagent containing malachite green, followed by incubation for 30 min at room temperature. The optical density at 620 nm was measured by using a microplate reader. ATP/GTPase activity was calculated by comparison to a phosphate standard curve.

### Cryo-EM

Purified wt hSAMD9L or **Δ**SAM-FL hSAMD9L (3.5 μL) was applied to glow-discharged UltraAuFoil R1.2/1.3 300 mesh gold grids (Electron Microscopy Sciences). Plunge freezing was performed by using a Vitrobot Mark IV (Thermo Fisher Scientific) with a blot force of 4 and blot time of 4.5 s at 85% humidity and 10 °C. Cryo-EM movies were acquired on a Talos Arctica transmission electron microscope (Thermo Fisher Scientific) operating at 200 kV, equipped with a K3 direct electron detector. Data were collected at a nominal magnification of 130,000×, corresponding to a pixel size of 0.63 Å. Automated data acquisition was carried out by using EPU software, with a defocus range of −0.5 μm to −2.5 μm. The total electron dose was 53.7 e⁻/Å² for wt hSAMD9L and 60.3 e⁻/Å² for **Δ**SAM-FL hSAMD9L.

### Data processing for Full-length wt hSAMD9L and **Δ**SAM-FL hSAMD9L

Details of cryo-EM data processing are provided in Suppl Figures 19 and 20; the overall steps are summarized below. For wt hSAMD9L, all movies were processed by using cryoSPARC v4.6 (48). Initial steps included patch motion correction and patch CTF estimation, followed by automated blob-based particle picking, which yielded 2,608,500 particles from 11,304 micrographs. Particles were extracted by using a box size of 512 pixels. After four rounds of 2D classification, ab-initio reconstruction, and non-uniform refinement, a reasonable map was obtained and used to generate templates for further particle picking. Template-based picking resulted in 7,360,068 particles. After three additional rounds of 2D classification, ab initio reconstruction, heterogeneous refinement, and non-uniform refinement, a consensus map was obtained at a resolution of 2.96 Å. To distinguish monomeric and dimeric forms, 3D classification was performed. This separated the monomer population and improved the monomer map resolution to 2.84 Å. Further 3D classification and refinement of the dimeric particles resulted in a final map at 2.89 Å resolution. The processed cryo-EM map was validated with the 3DFSC server (49).

In the case of **Δ**SAM-FL hSAMD9L, a total of 1,996,453 particles were picked from 10,575 micrographs. Particles were extracted by using a box size of 512 pixels. After three rounds of 2D classification, ab-initio reconstruction, and non-uniform refinement, an initial map was generated and used to create templates for further particle picking. Template-based picking yielded 4,277,132 particles. These were subjected to three additional rounds of 2D classification, ab-initio reconstruction, heterogeneous refinement, and non-uniform refinement, resulting in separate monomeric and dimeric reconstructions. Final maps were obtained at resolutions of 3.96 Å for the monomer and 3.67 Å for the dimer, respectively. The processed cryo-EM map was validated with 3DFSC server (49).

### Model building

The initial model of hSAMD9L was generated by using AlphaFold (50). The predicted structure was manually truncated to remove the SAM domain, the flexible linker region, and the AlbA domain. This truncated model was fitted into the monomeric cryo-EM map by using ChimeraX-1.10.1 (51) and rebuilt by manual tracing in Coot-1.0 (52). For the dimer model, individual domains were manually adjusted to fit the conformationally distinct protomer. Real-space refinement was carried out by using Phenix-1.21.2 (53), and the final models were validated using MolProbity (54). All structural figures were generated by using PyMOL-3.0.3 (55) and ChimeraX-1.10.1 (51).

## Data availability

The final coordinates and cryo-EM maps supporting the findings of this study have been deposited in the Worldwide Protein Data Bank (https://www.wwpdb.org) under the following accession codes: PDB IDs (9YVR and 9YWC for wt hSAMD9L monomer and dimer, respectively, and 9YWO and 9YWR for **Δ**SAM-FL hSAMD9L respectively) and EMDB IDs (EMD-73532 and EMD-73544 for wt hSAMD9L monomer and dimer, respectively, and EMD-73554 and EMD73557 for **Δ**SAM-FL hSAMD9L, respectively).

---

**Suppl figure 1.**
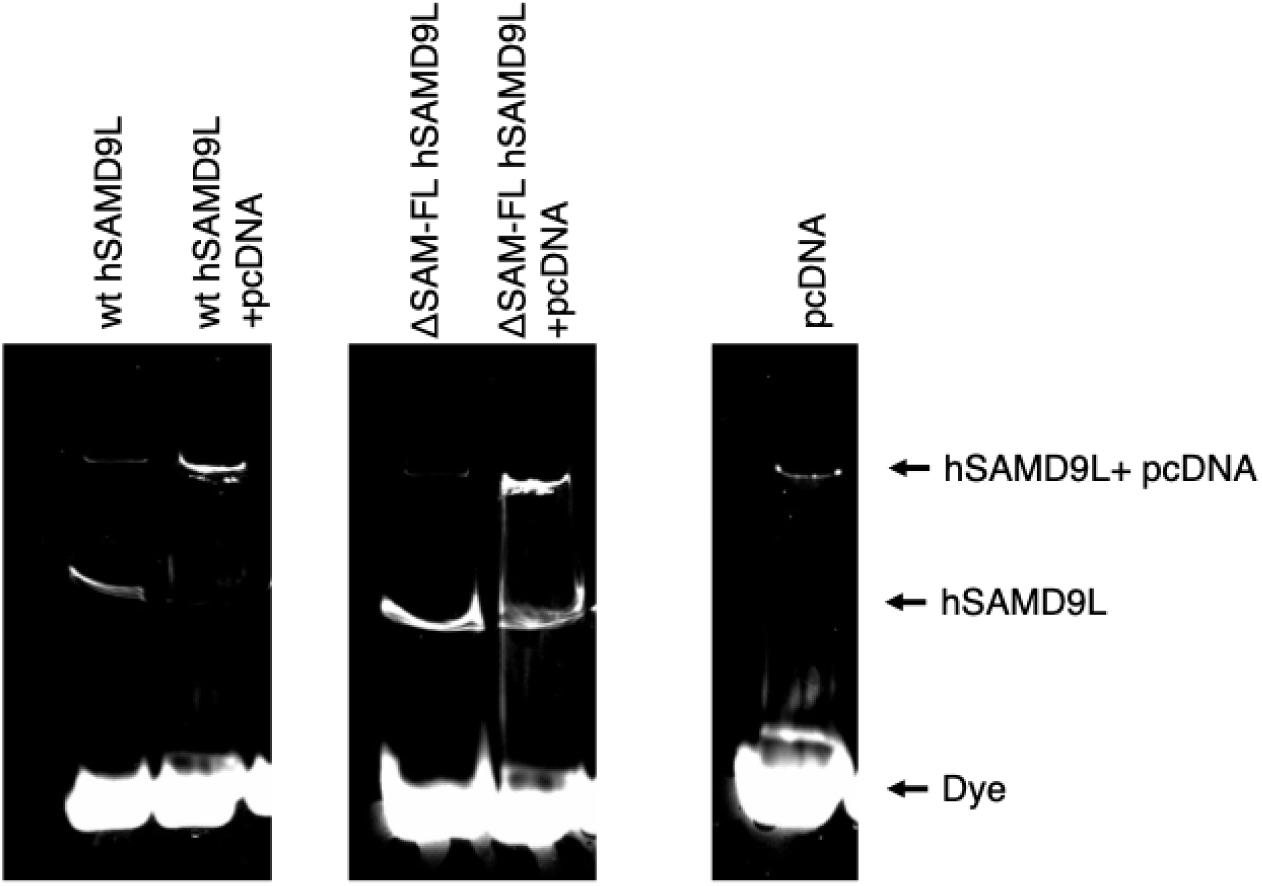
Native PAGE analysis of hSAMD9L oligomerization upon dsDNA binding. Circular plasmid DNA (pcDNA) was incubated with wt hSAMD9L or ΔSAM-FL hSAMD9L followed by native polyacrylamide gel electrophoresis. Gels were imaged under UV illumination. The middle arrowhead marks free hSAMD9L, and the upper arrowhead marks DNA–protein complexes.

**Suppl figure 2.**
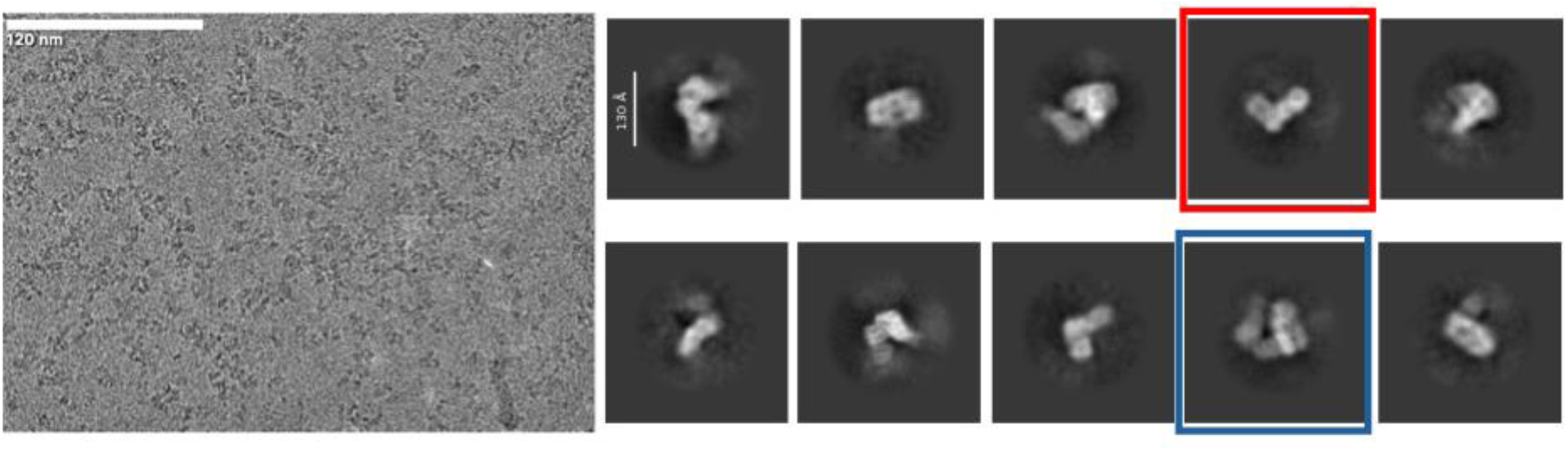
wt hSAMD9L Cryo-EM data collection and processing. **Left**: Representative micrograph acquired during data collection. **Right**: Examples of 2D classification results. Representative 2D classes of monomer and dimer, outlined in red and blue, respectively.

**Suppl figure 3.**
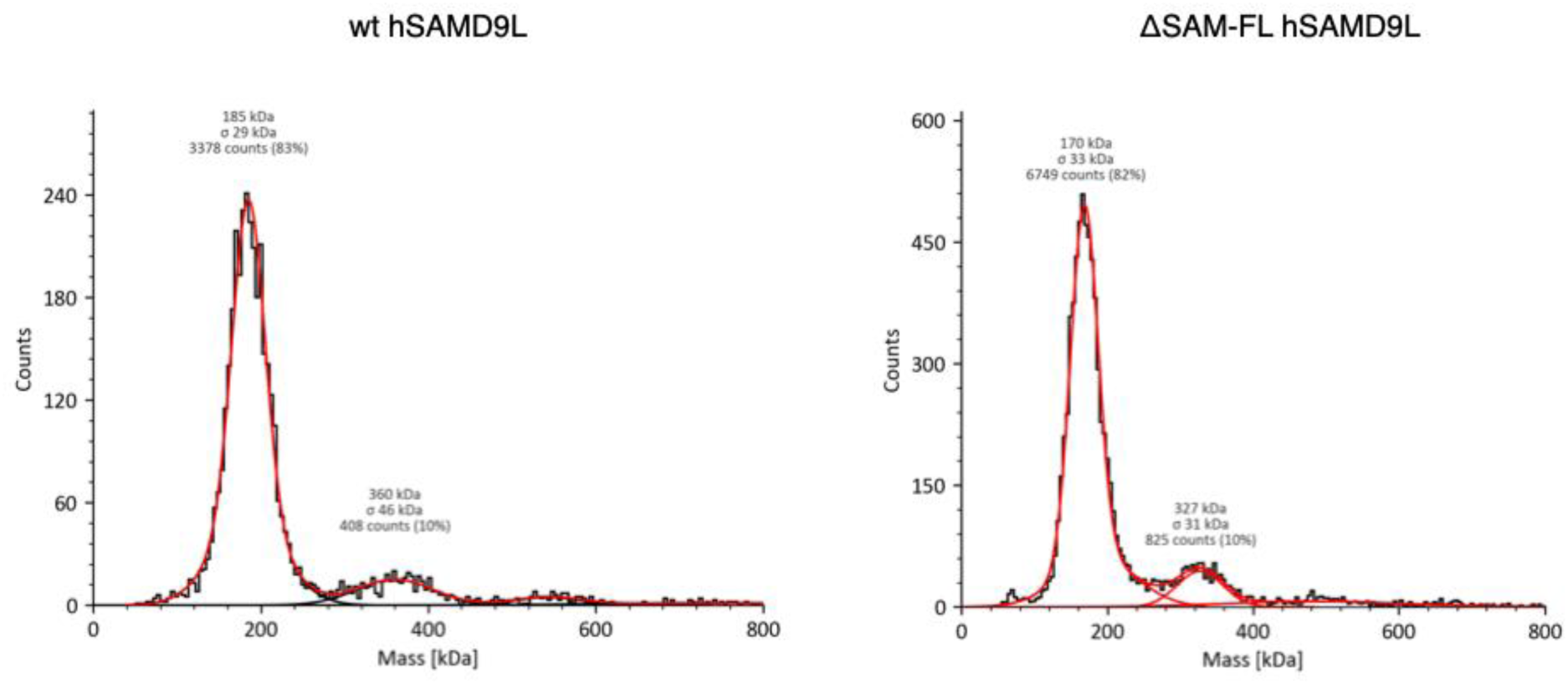
Mass photometry analysis of wt and ΔSAM-FL hSAMD9L. **Left**: wt hSAMD9L displays two major species with estimated molecular masses of 185 kDa (SD = 29) and 360 kDa (SD = 46), consistent with the theoretical masses of the monomer (188.6 kDa) and dimer (377.2 kDa), respectively. **Right**: ΔSAM-FL hSAMD9L shows molecular mass estimates of 170 kDa (SD = 33) and 327 kDa (SD = 31), in agreement with the theoretical monomer and dimer masses of 167.3 kDa and 334.6 kDa, respectively.

**Suppl figure 4.**
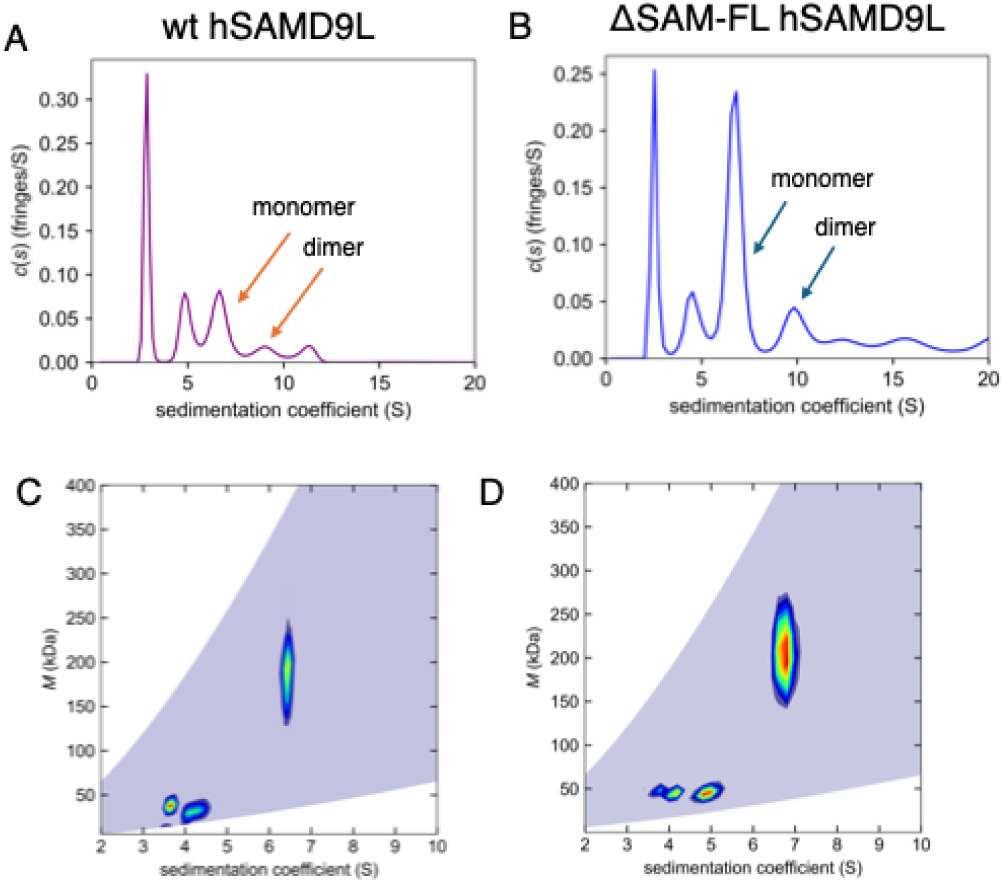
Sedimentation velocity AUC wt hSAMD9L and ΔSAM-FL hSAMD9L. The sedimentation velocity profiles (fringe displacement) were fitted to a continuous sedimentation coefficient distribution model *c(s)*, as well as to a 2D size-and-shape distribution model, *c(s,f/f_0_)*. (**A**) Single *c(s)* distribution of wt hSAMD9L at 0.47 *µM*. (**B**) Single *c(s)* distribution of ΔSAM-FL hSAMD9L at 1.01 *µM*. (**C)** Velocity data of wt hSAMD9L analyzed with the *c(s,f/f_0_)* model and transformed to a *c(s,M)* contour plot as a heat map with increasing color temperature to maximum fringes/*S* value in red. (**D**) Velocity data of ΔSAM-FL hSAMD9L analyzed with the *c(s,f/f_0_)* model and transformed to a *c(s,M)* contour plot as a heat map with increasing color temperature to maximum fringes/*S* value in red. Experiments were conducted in 50 mM HEPES-NaOH pH 7.4, 300 mM NaCl buffer at 20°C at rotor speed of 42,000 rpm. The *s*- and mass-values are listed in tables.

**Suppl figure 5.**
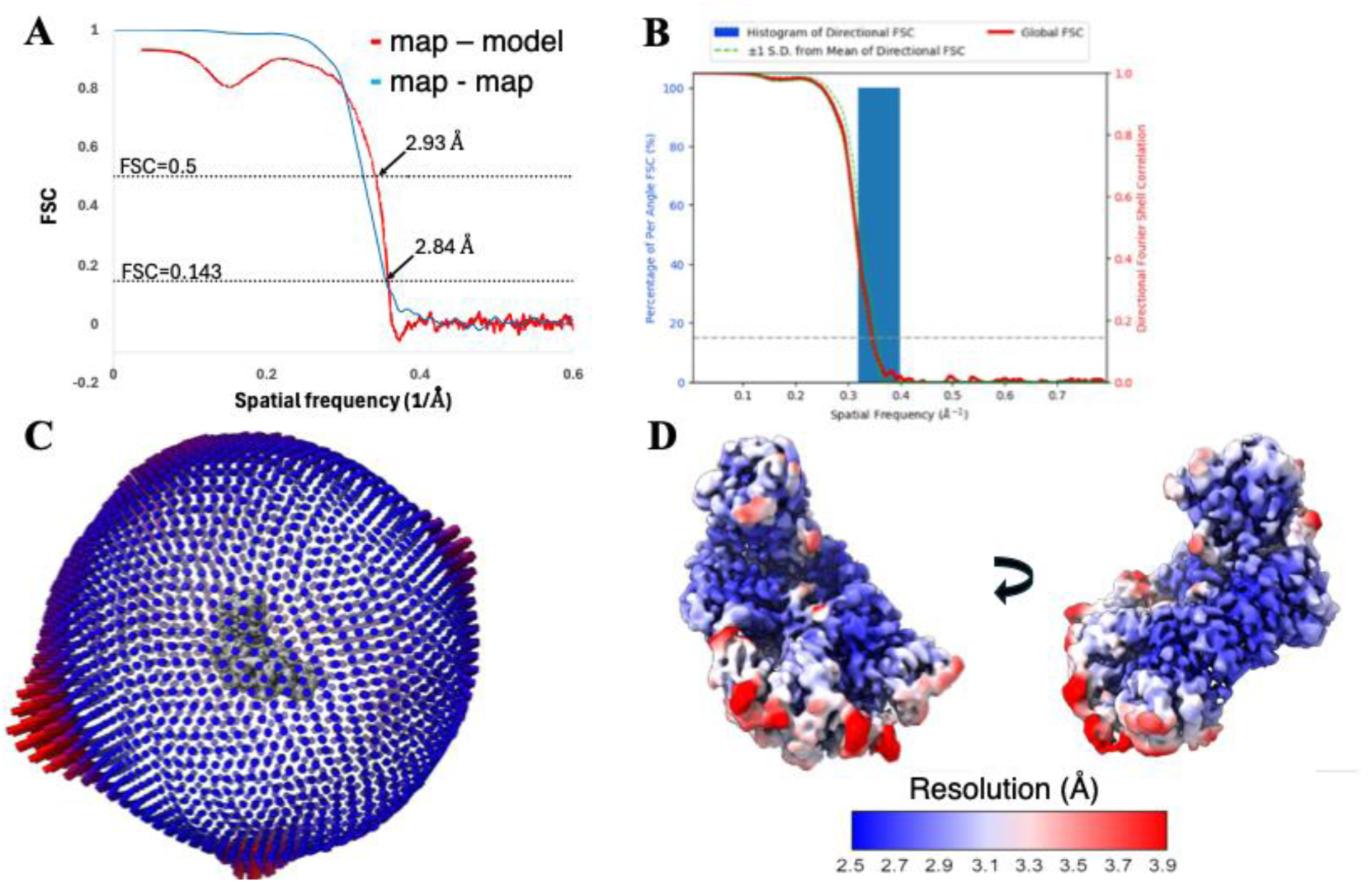
cryo-EM map and model quality assessment of the wt hSAMD9L monomer. A: Fourier shell correlation (FSC) curves plotted against spatial frequency (1/Å). The blue line represents the FSC between two half maps, and the red line shows the FSC between the refined model and the full map. B: 3D Fourier Shell Correlation (3DFSC) plot for the final cryo-EM reconstruction. C: Particle orientation distribution for the final cryo-EM map. D: Local resolution map of the final cryo-EM reconstruction, color-coded by resolution.

**Suppl figure 6.**
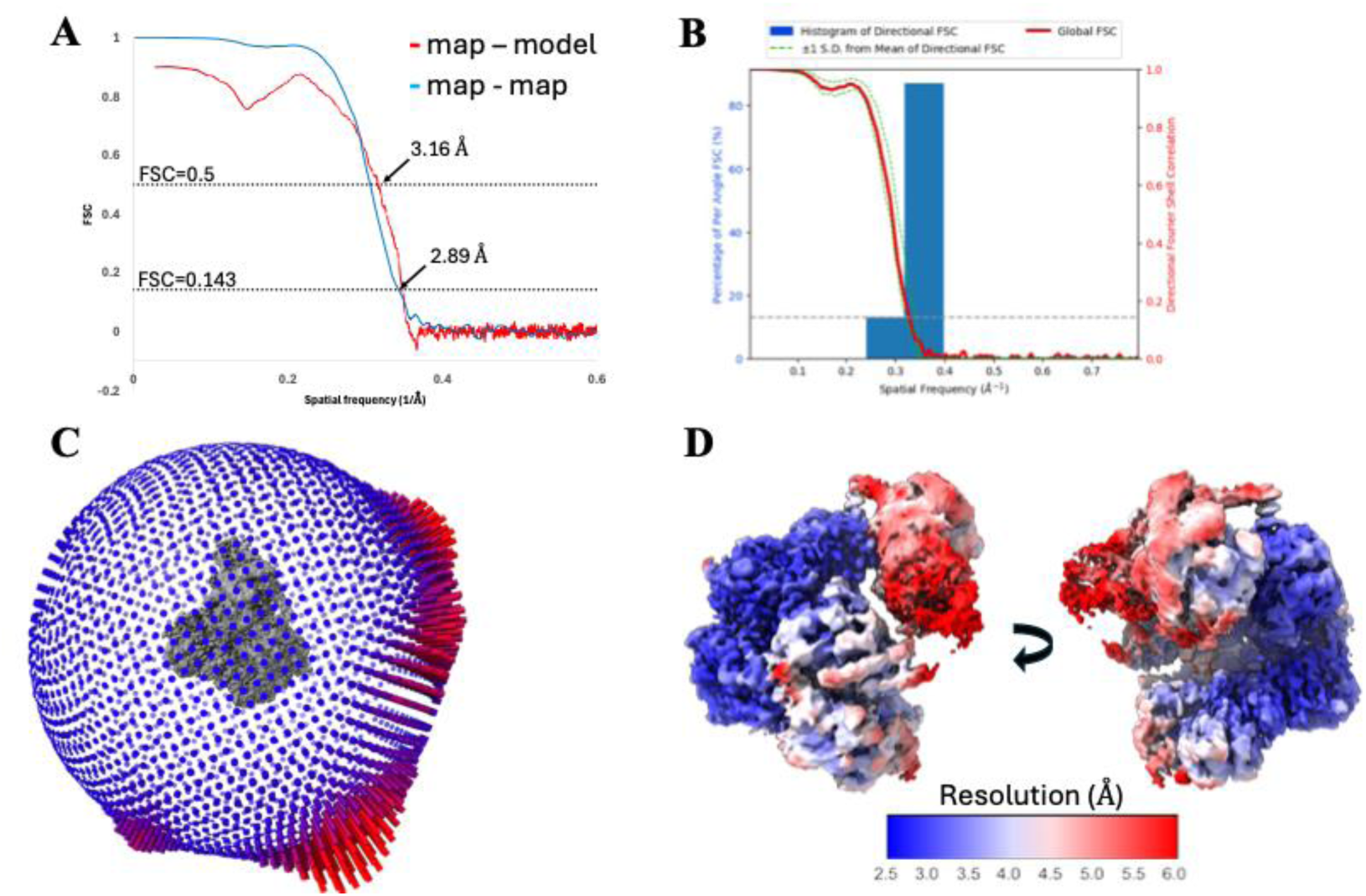
Cryo-EM map and model quality assessment of the wt hSAMD9L dimer. A: Fourier shell correlation (FSC) curves plotted against spatial frequency (1/Å). The blue line represents the FSC between two half maps, and the red line shows the FSC between the refined model and the full map. B: 3D Fourier Shell Correlation (3DFSC) plot for the final cryo-EM reconstruction. C: Particle orientation distribution for the final cryo-EM map. D: Local resolution map of the final cryo-EM reconstruction, color-coded by resolution.

**Suppl figure 7.**
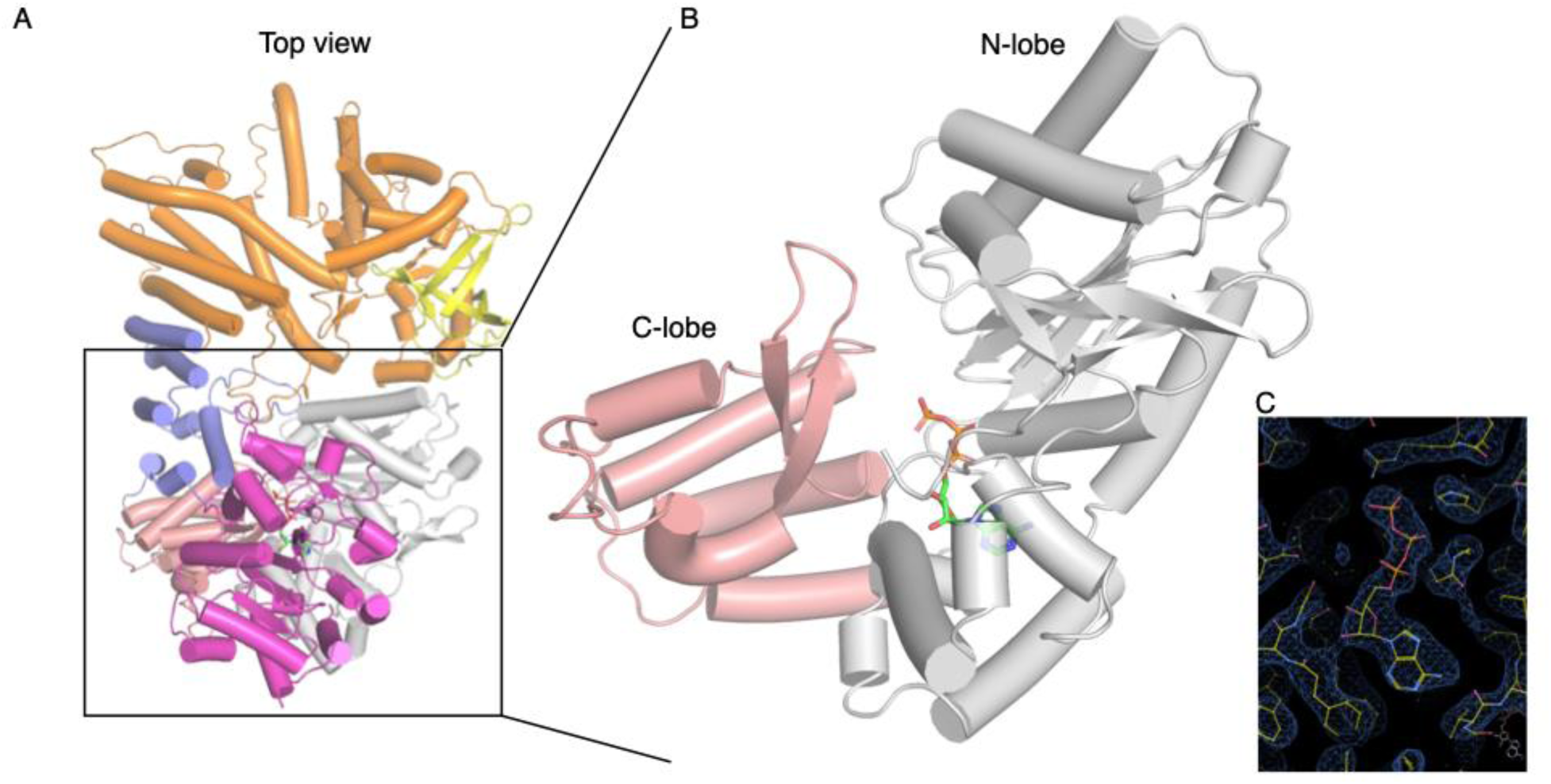
ATP assignment in the N-lobe of the NTPase domain. The cryo-EM map of the N-lobe of the NTPase domain with assigned bound ATP. **A**: Top view of wt hSAMD9L. **B**: Focused view of the NTPase domain. **C**: Zoomed-in cryo-EM map of the NTPase domain and ATP.

**Suppl figure 8.**
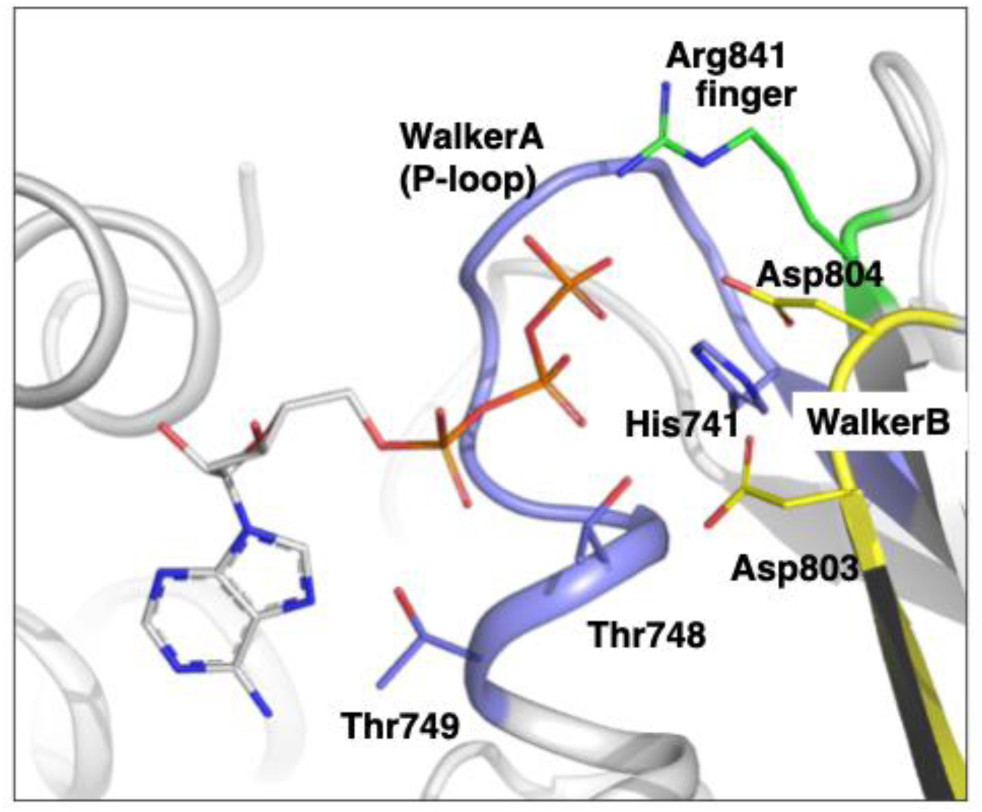
Detailed view of the NTP-binding site in the N-lobe of the NTPase domain in wt hSAMD9L. Key conserved motifs involved in nucleotide binding are highlighted: **Blue** – Walker A (P-loop), **Yellow** – Walker B, and **Green** – the “finger” arginine residue.

**Suppl figure 9.**
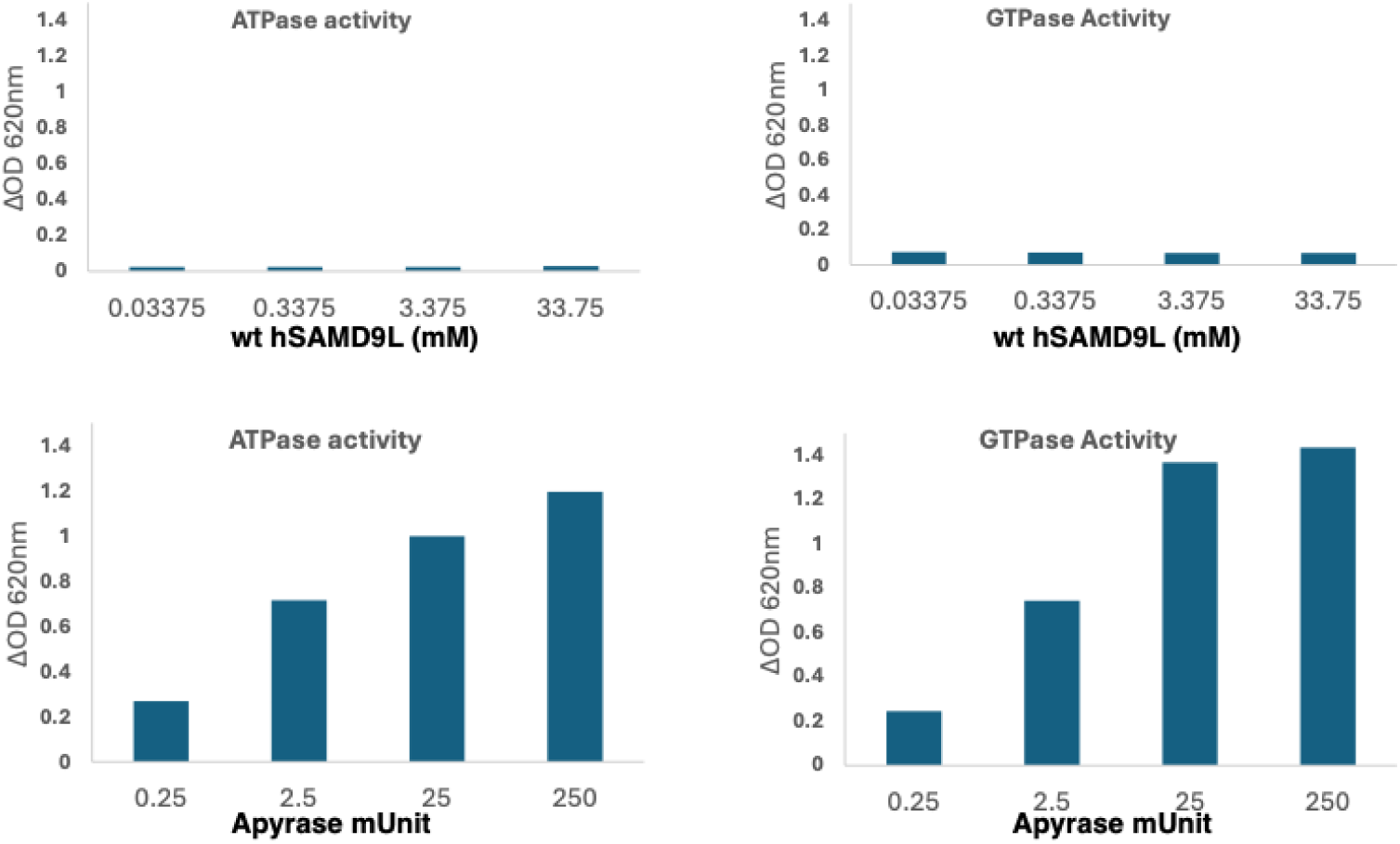
Evaluation of ATPase and GTPase activity of wt hSAMD9L. **Top**: wt hSAMD9L did not exhibit detectable ATPase or GTPase activity across a range of protein concentrations. **Left**: ATP as substrate. **Right**: GTP as substrate. **Bottom**: Apyrase used as a positive control for ATPase and GTPase activity assays.

**Suppl figure 10.**
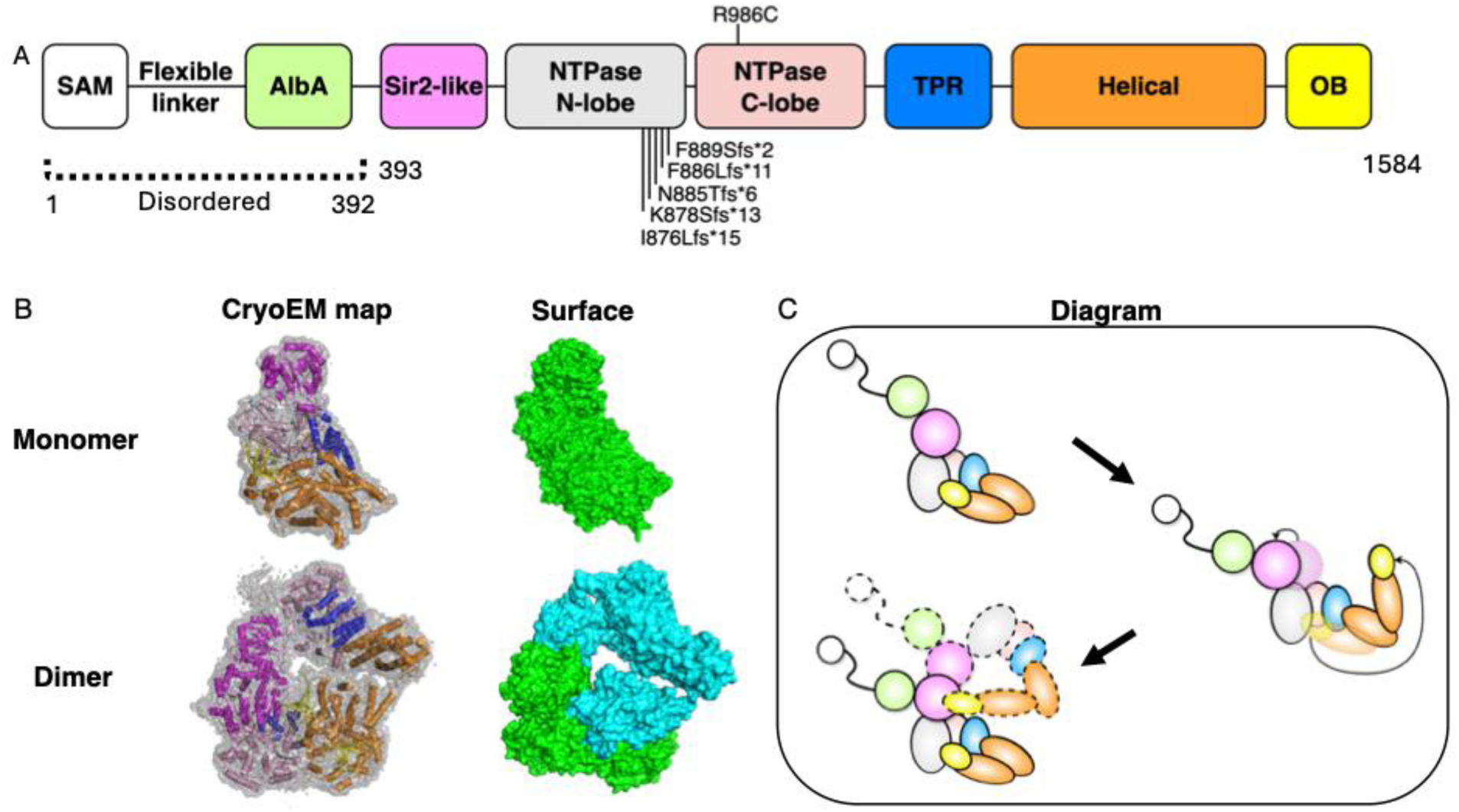
cryo-EM structure of the dimer of wt hSAMD9L. **A: Domain organization of wt hSAMD9L.** Schematic diagram of the full-length structure showing annotated domains. The region from the SAM to the AlbaA domain is not resolved in the map. The NTPase domain is divided into N-lobe and C-lobe for clarity. Disease-associated missense mutation and frameshift mutations are mapped to the NTPase domain. The region between the TPR and OB domains is designated as the “helical domain.” **B: Cryo-EM map and structural model. Left**: Cryo-EM maps of the wt hSAMD9L monomer and dimer are shown, with the fitted atomic models represented as ribbon diagrams. **Right**: Surface representations of the atomic models for both monomer and dimer. **C: Schematic representation of conformational changes associated with dimer formation.** A proposed model illustrating the conformational transition of wt hSAMD9L upon dimerization. The top-left panel shows the monomeric form; domain opening is illustrated in the right panel. The bottom-left panel depicts the docking of the open form to an unaltered monomer to form the dimer.

**Suppl figure 11.**
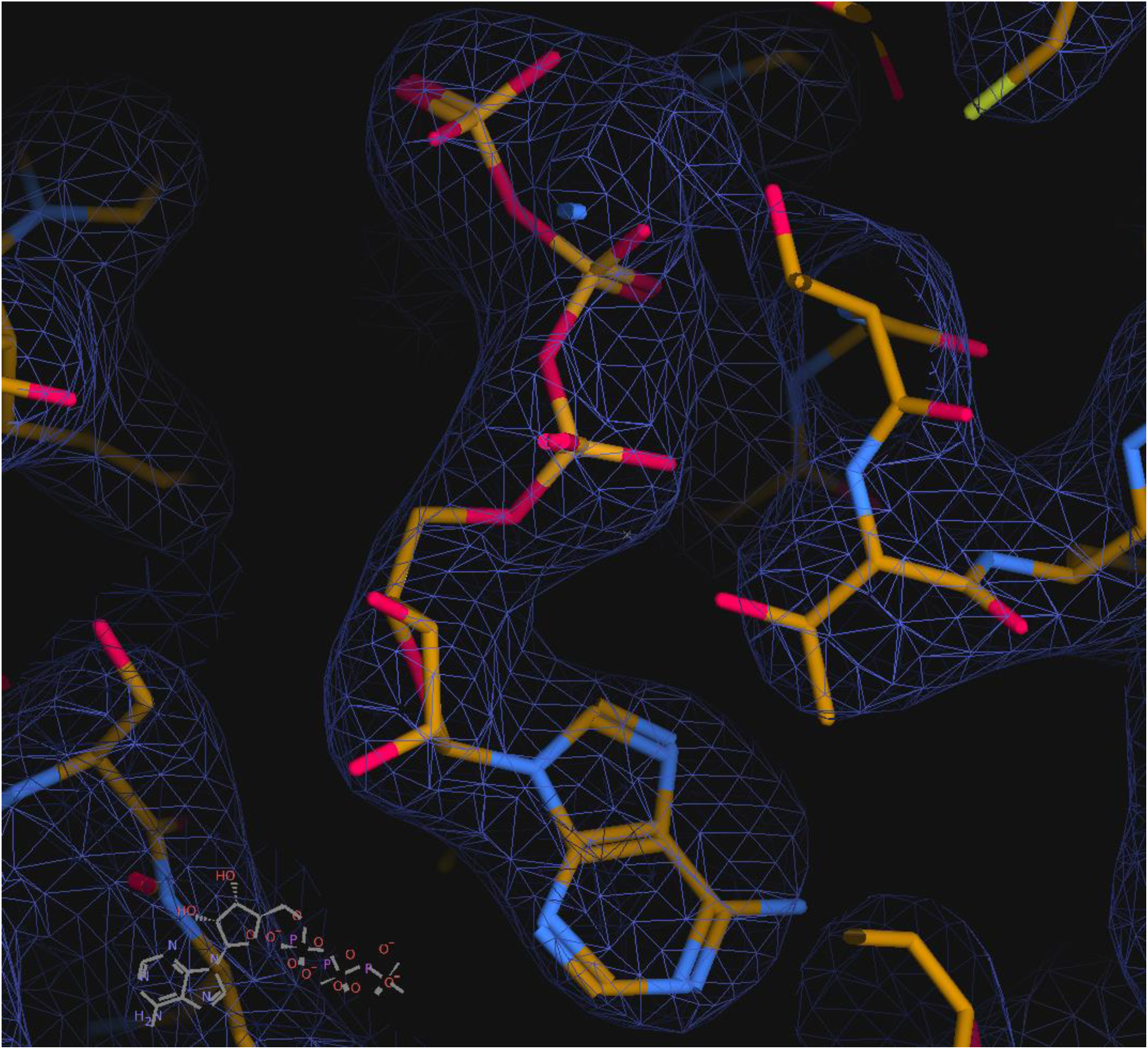
cryo-EM map for NTP in wt hSAMD9L dimer structure. Cryo-EM density observed in the NTPase domain of the protomer A in the dimeric wt hSAMD9L structure.

**Suppl figure 12.**
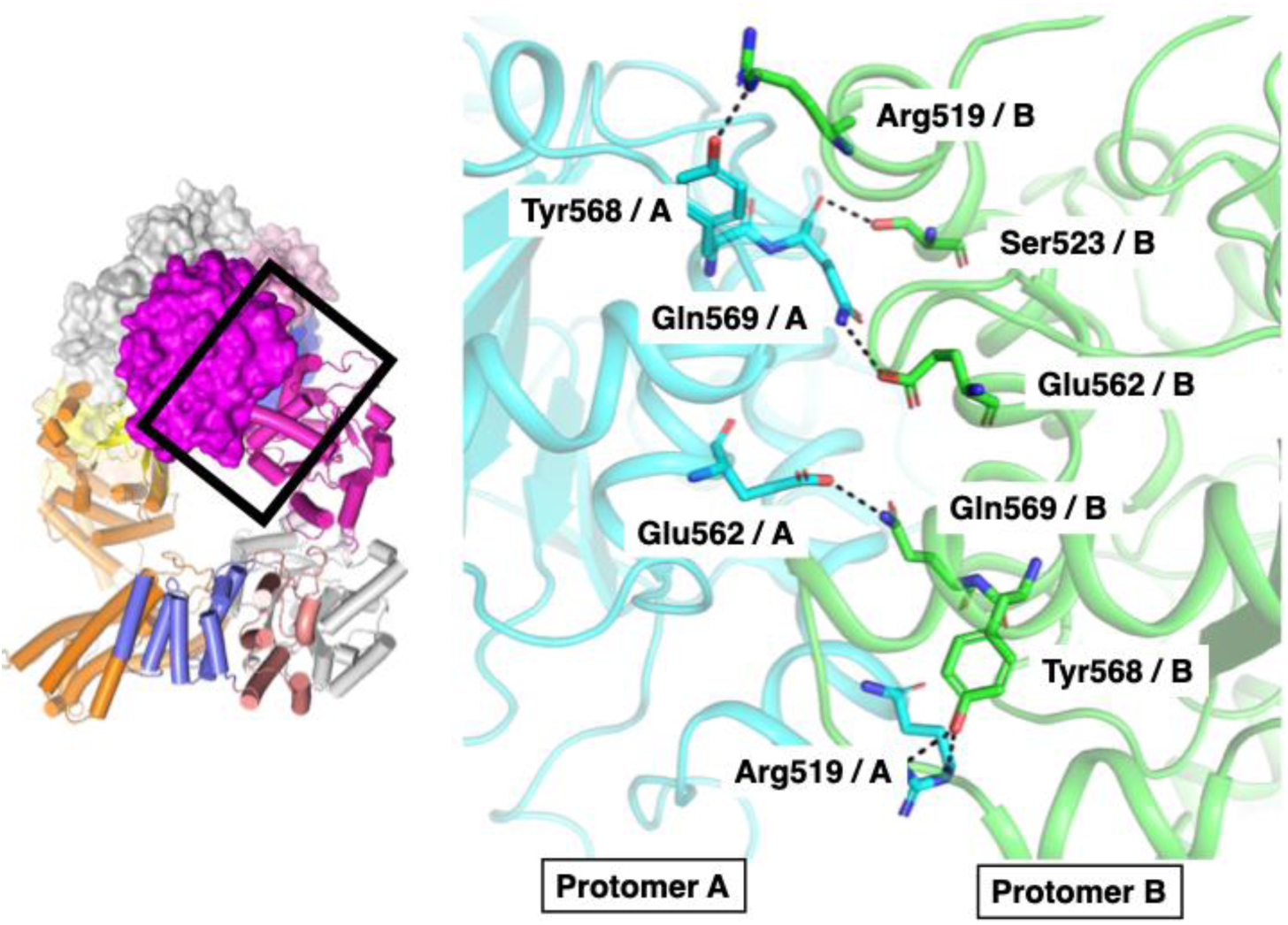
Sir2–Sir2 interaction interface in the dimeric structure of wt hSAMD9L. The two protomers are shown as ribbon models, with Protomer A in cyan and Protomer B in green. Residues involved in the Sir2–Sir2 interaction are displayed as sticks.

**Suppl figure 13.**
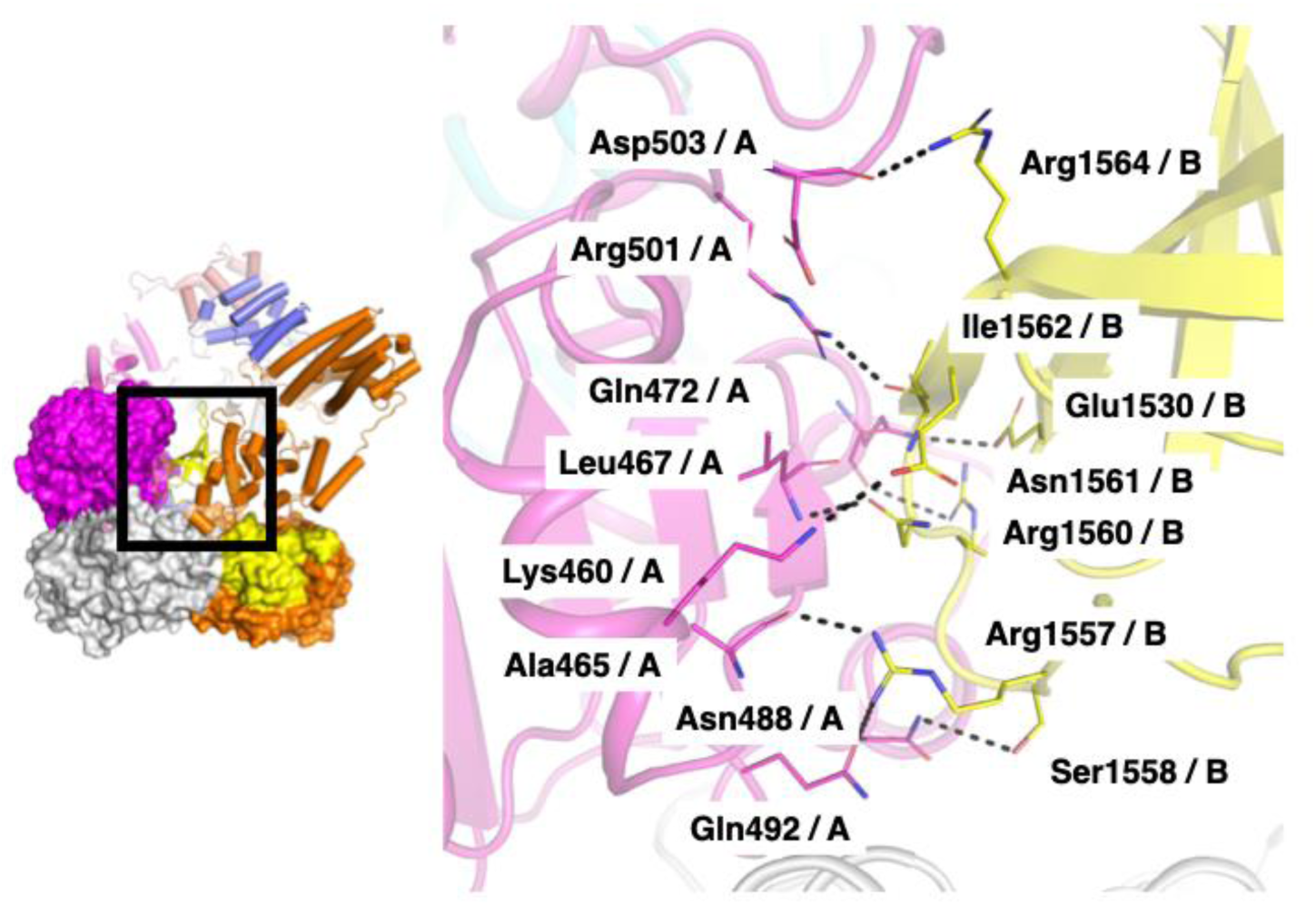
Sir2–OB interaction interface in the dimeric structure of wt hSAMD9L. The two protomers are shown as ribbon models, with Protomer A in purple and Protomer B in yellow. Residues involved in the Sir2–OB interaction are displayed as sticks.

**Suppl figure 14.**
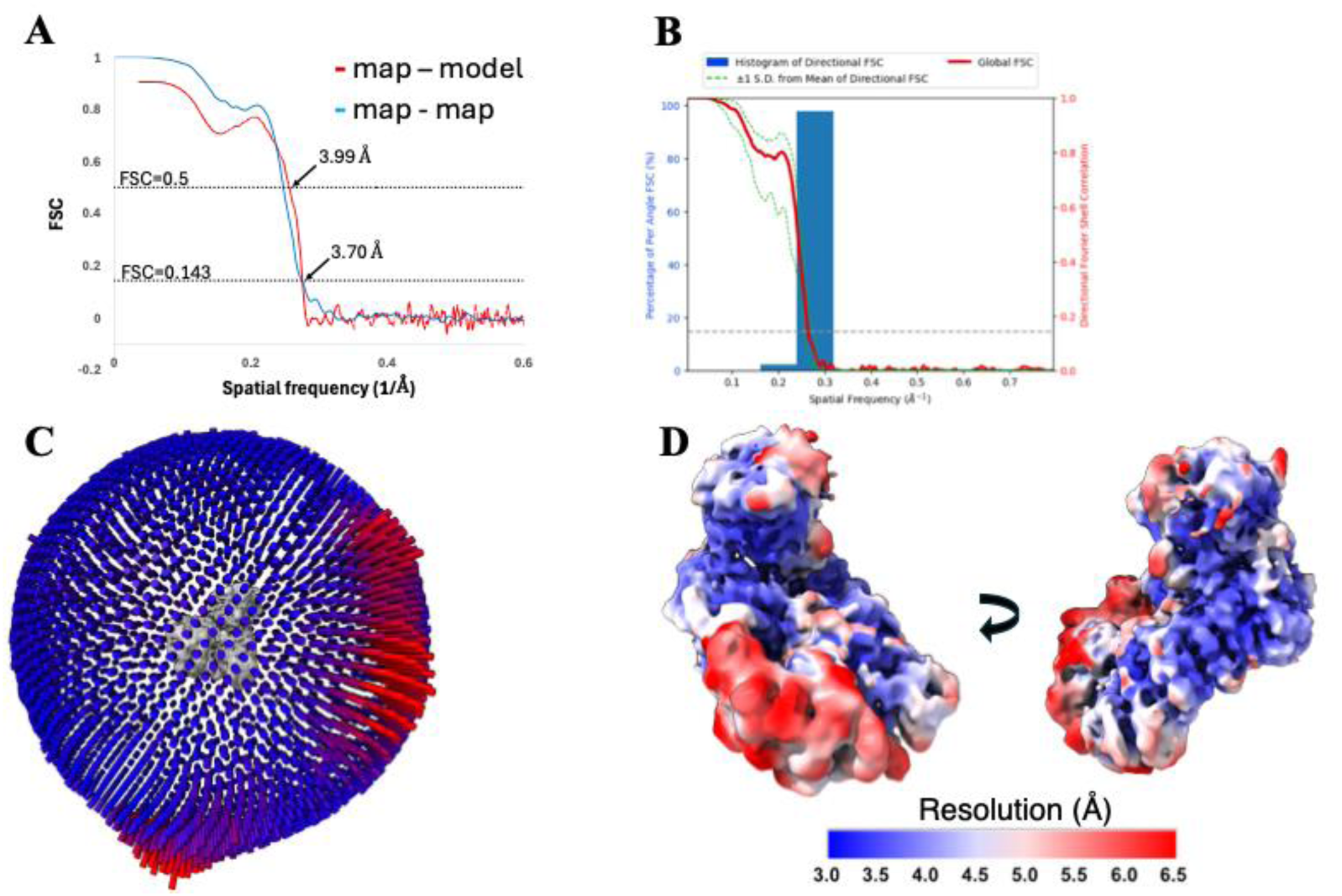
cryo-EM map and model quality assessment of the ΔSAM-FL hSAMD9L monomer. A: Fourier shell correlation (FSC) curves plotted against spatial frequency (1/Å). The blue line represents the FSC between two half maps, and the red line shows the FSC between the refined model and the full map. B: 3D Fourier Shell Correlation (3DFSC) plot for the final cryo-EM reconstruction. C: Particle orientation distribution for the final cryo-EM map. D: Local resolution map of the final cryo-EM reconstruction, color-coded by resolution.

**Suppl figure 15.**
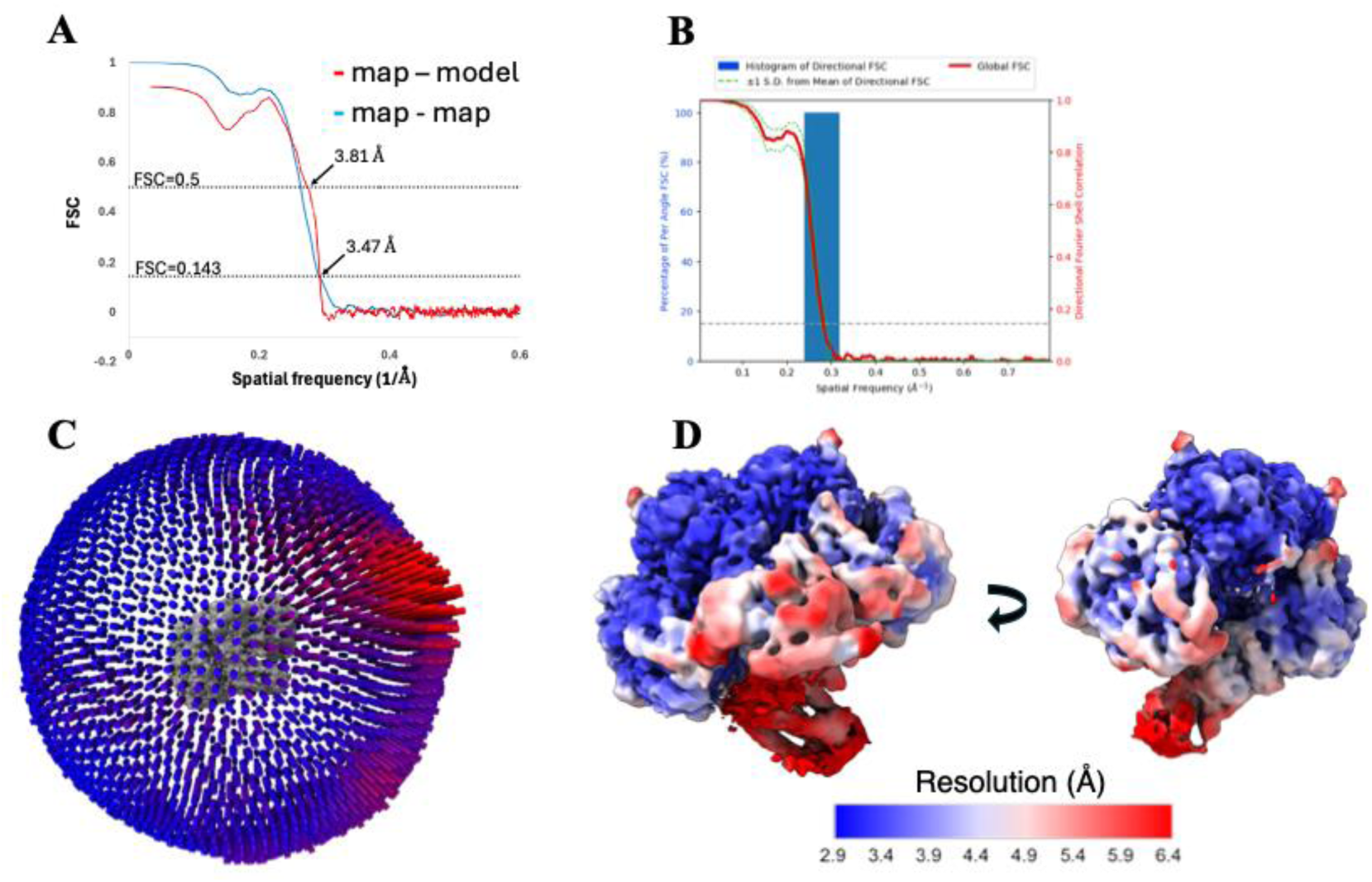
cryo-EM map and model quality assessment of the ΔSAM-FL hSAMD9L dimer. A: Fourier shell correlation (FSC) curves plotted against spatial frequency (1/Å). The blue line represents the FSC between two half maps, and the red line shows the FSC between the refined model and the full map. B: 3D Fourier Shell Correlation (3DFSC) plot for the final cryo-EM reconstruction. C: Particle orientation distribution for the final cryo-EM map. D: Local resolution map of the final cryo-EM reconstruction, color-coded by resolution.

**Suppl figure 16.**
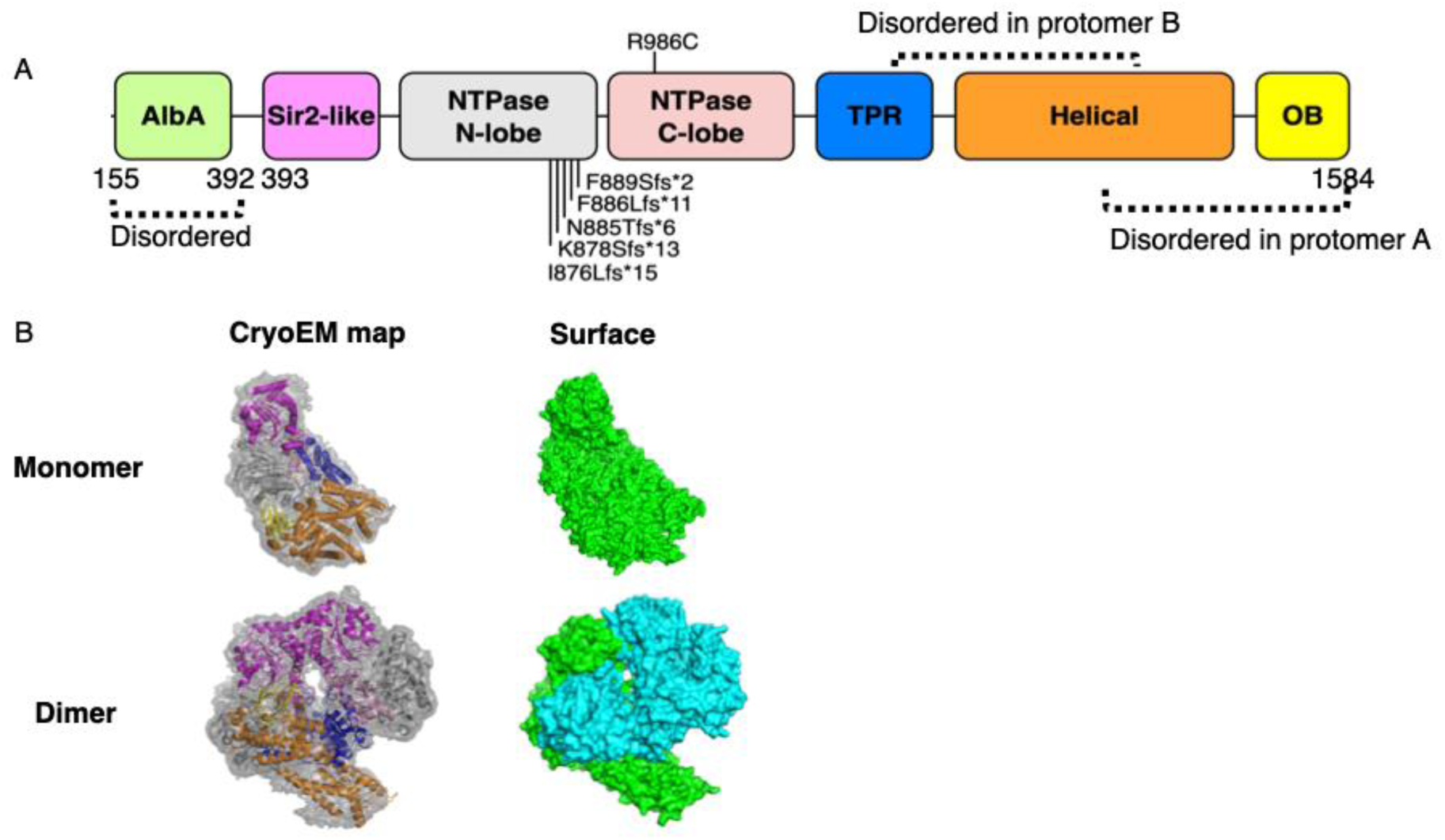
cryo-EM structure of ΔSAM-FL hSAMD9L. **A: Domain organization of hSAMD9L.** Schematic diagram of the structure showing annotated domains. The AlbA domain and part of Helical domain and OB domain are not resolved in the map. The NTPase domain is divided into N-lobe and C-lobe for clarity. Disease-associated missense mutation and frameshift mutations are mapped to the NTPase domain. **B: Cryo-EM map and structural model. Left**: Cryo-EM density maps of the ΔSAM-FL hSAMD9L monomer and dimer are shown, with the fitted atomic models represented as ribbon diagrams. **Right**: Surface representations of the atomic models for both monomer and dimer.

**Suppl figure 17.**
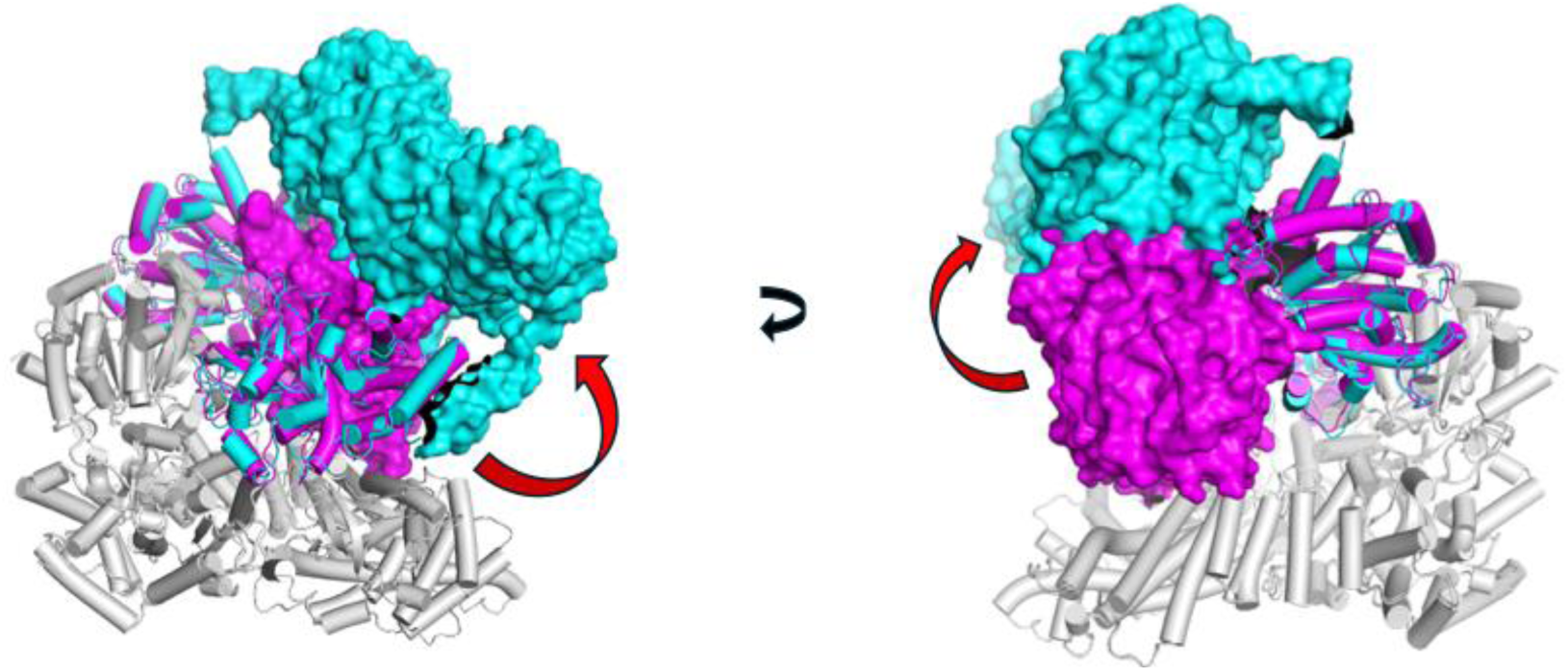
Superimposed structures of wt hSAMD9L (cyan) and ΔSAM-FL hSAMD9L (purple). Protomer A from both constructs is shown as a white cylinder representation. In Protomer B, the SAM domain, a portion of the helical domain, and the OB domain are also displayed as cylinders. The NTPase, TPR, and part of the helical domains in Protomer B are shown in surface representation to highlight structural differences.

**Suppl figure 18.**
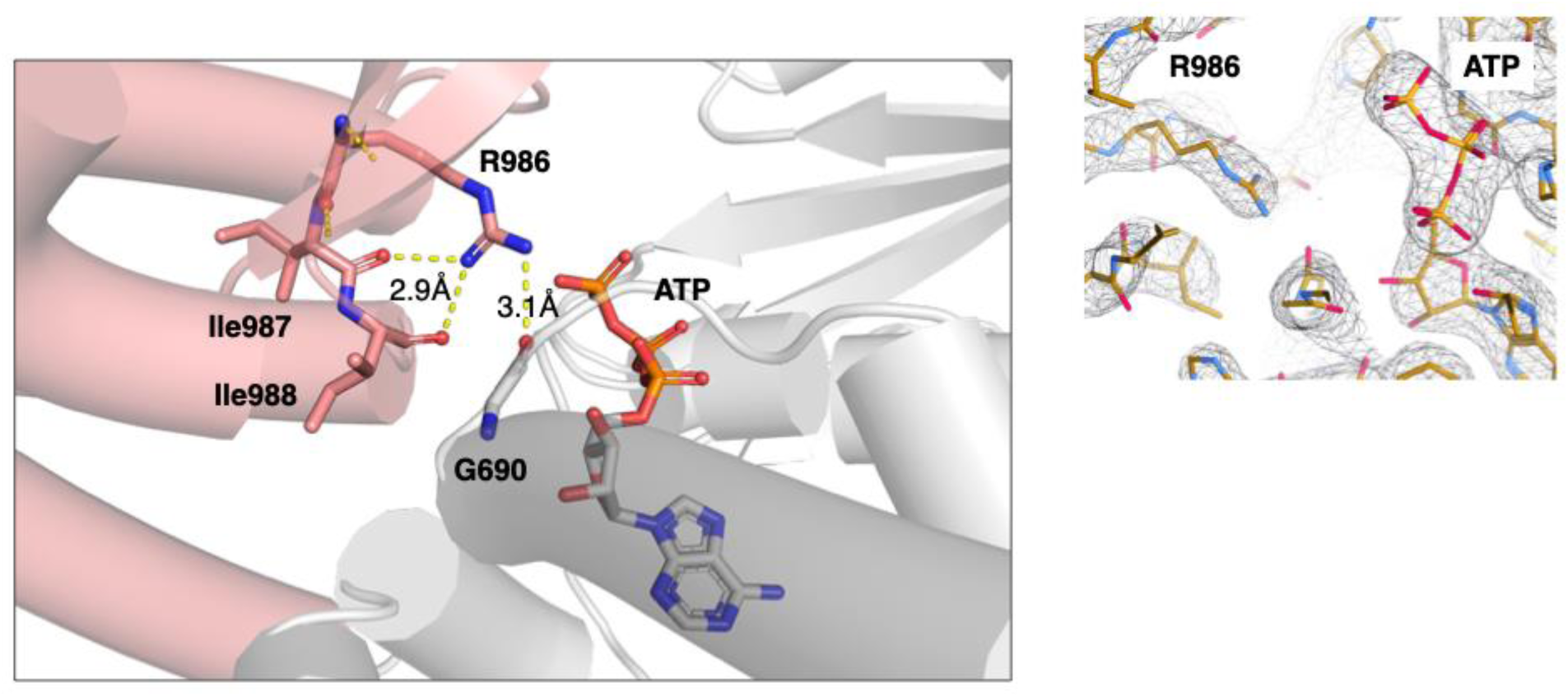
Structural context of the disease-associated R986C mutation in wt hSAMD9L. R986, a residue mutated in the disease-associated R986C variant, is in the C-lobe of the NTPase domain, positioned opposite the N-lobe. Structural analysis indicates that R986 engages in intramolecular interactions within the C-lobe. **Right**: Cryo-EM map of R986 and ATP visualized in *Coot*.

**Suppl figure 19.**
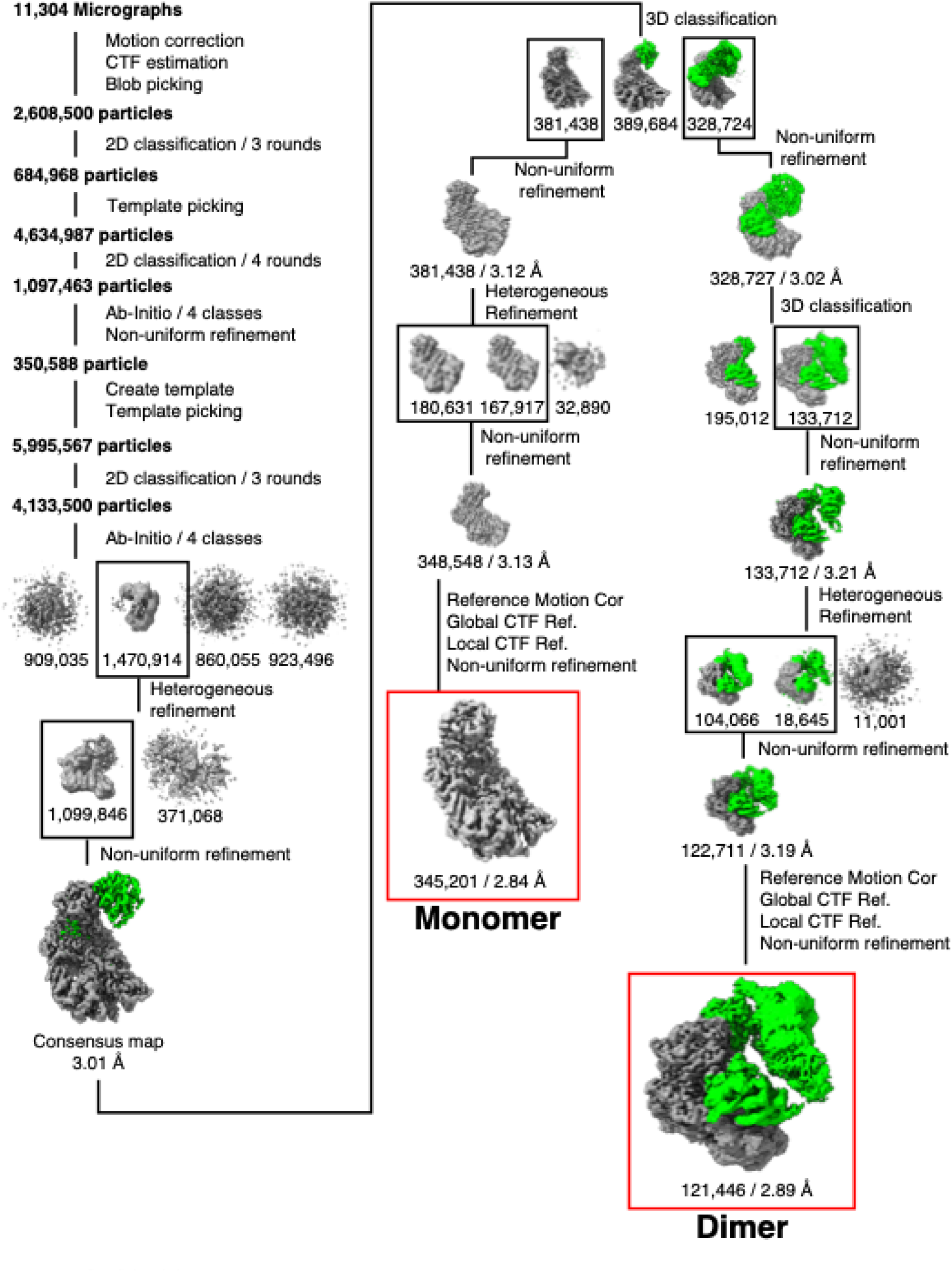
cryo-EM processing for wt hSAMD9L.

**Suppl figure 20.**
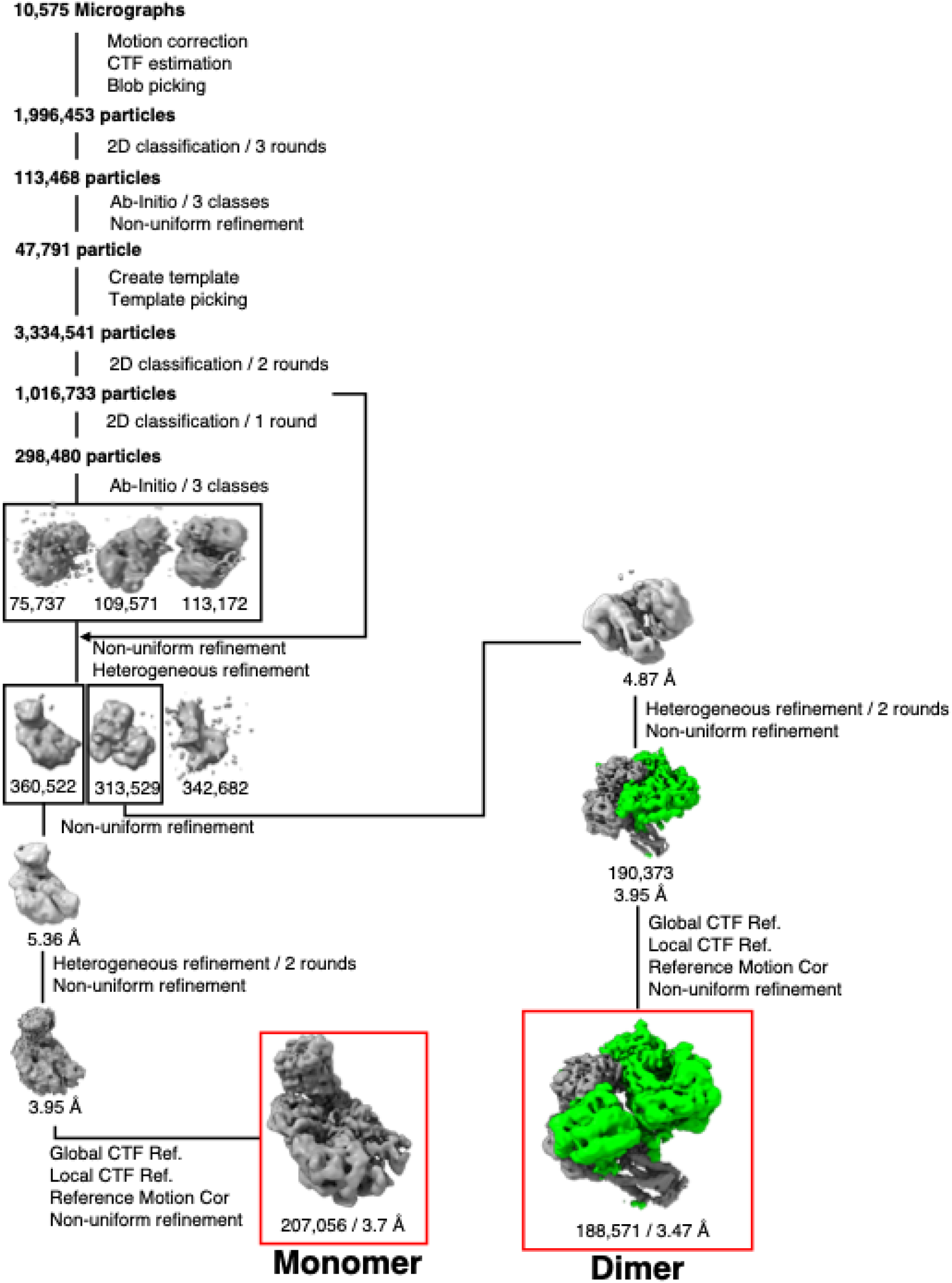
cryo-EM processing for ΔSAM-FL hSAMD9L.

## Notes

### Competing Interest Statement

The authors have declared no competing interest.

## References

1. Li CF, MacDonald JR, Wei RY, Ray J, Lau K, Kandel C, et al. Human sterile alpha motif domain 9, a novel gene identified as down-regulated in aggressive fibromatosis, is absent in the mouse. BMC Genomics. 2007;8:92.

2. McSweeney K, Hoover P, Ramirez-Solano M, Liu Q, Schwartz JR. Overexpression of human SAMD9 inhibits protein translation and alters MYC signaling resulting in cell cycle arrest. Exp Hematol. 2024;137:104249.

3. Xiang Y. Poxvirus Host-Range Determinants: SAMD9/9L and Beyond. Annu Rev Virol. 2025.

4. Liu J, McFadden G. SAMD9 is an innate antiviral host factor with stress response properties that can be antagonized by poxviruses. J Virol. 2015;89(3):1925–31.

5. Tesi B, Davidsson J, Voss M, Rahikkala E, Holmes TD, Chiang SCC, et al. Gain-of-function SAMD9L mutations cause a syndrome of cytopenia, immunodeficiency, MDS, and neurological symptoms. Blood. 2017;129(16):2266–79.

6. Legrand A, Dahoui C, De La Myre Mory C, Noy K, Guiguettaz L, Versapuech M, et al. SAMD9L acts as an antiviral factor against HIV-1 and primate lentiviruses by restricting viral and cellular translation. PLoS Biol. 2024;22(7):e3002696.

7. Maleknia S, Tavassolifar MJ, Mottaghitalab F, Zali MR, Meyfour A. Identifying novel host-based diagnostic biomarker panels for COVID-19: a whole-blood/nasopharyngeal transcriptome meta-analysis. Mol Med. 2022;28(1):86.

8. Hou G, Beatty W, Ren L, Ooi YS, Son J, Zhu Y, et al. SAMD9 senses cytosolic double-stranded nucleic acids in epithelial and mesenchymal cells to induce antiviral immunity. Nat Commun. 2025;16(1):3756.

9. Chen DH, Below JE, Shimamura A, Keel SB, Matsushita M, Wolff J, et al. Ataxia-Pancytopenia Syndrome Is Caused by Missense Mutations in SAMD9L. Am J Hum Genet. 2016;98(6):1146–58.

10. Meng X, Xiang Y. RNA granules associated with SAMD9-mediated poxvirus restriction are similar to antiviral granules in composition but do not require TIA1 for poxvirus restriction. Virology. 2019;529:16–22.

11. Sivan G, Glushakow-Smith SG, Katsafanas GC, Americo JL, Moss B. Human Host Range Restriction of the Vaccinia Virus C7/K1 Double Deletion Mutant Is Mediated by an Atypical Mode of Translation Inhibition. J Virol. 2018;92(23).

12. Zhang F, Ji Q, Chaturvedi J, Morales M, Mao Y, Meng X, et al. Human SAMD9 is a poxvirus-activatable anticodon nuclease inhibiting codon-specific protein synthesis. Sci Adv. 2023;9(23):eadh8502.

13. Jagadeesh J, Vembar SS. Evolution of sequence, structural and functional diversity of the ubiquitous DNA/RNA-binding Alba domain. Sci Rep. 2024;14(1):30363.

14. Zhang LK, Chai F, Li HY, Xiao G, Guo L. Identification of host proteins involved in Japanese encephalitis virus infection by quantitative proteomics analysis. J Proteome Res. 2013;12(6):2666–78.

15. Gahr S, Perinetti Casoni G, Falk-Paulsen M, Maschkowitz G, Bryceson YT, Voss M. Viral host range factors antagonize pathogenic SAMD9 and SAMD9L variants. Exp Cell Res. 2023;425(2):113541.

16. Meng X, Zhang F, Yan B, Si C, Honda H, Nagamachi A, et al. A paralogous pair of mammalian host restriction factors form a critical host barrier against poxvirus infection. PLoS Pathog. 2018;14(2):e1006884.

17. Legrand A, Demeure R, Chantharath A, Rey C, Baltenneck J, Gilchrist CLM, et al. Evolutionary characterization of antiviral SAMD9/9L across kingdoms supports ancient convergence and lineage-specific adaptations. Nat Ecol Evol. 2025;9(12):2206–22.

18. Bluteau O, Sebert M, Leblanc T, Peffault de Latour R, Quentin S, Lainey E, et al. A landscape of germ line mutations in a cohort of inherited bone marrow failure patients. Blood. 2018;131(7):717–32.

19. Schwartz JR, Ma J, Lamprecht T, Walsh M, Wang S, Bryant V, et al. The genomic landscape of pediatric myelodysplastic syndromes. Nat Commun. 2017;8(1):1557.

20. Schwartz JR, Wang S, Ma J, Lamprecht T, Walsh M, Song G, et al. Germline SAMD9 mutation in siblings with monosomy 7 and myelodysplastic syndrome. Leukemia. 2017;31(8):1827–30.

21. Wong JC, Bryant V, Lamprecht T, Ma J, Walsh M, Schwartz J, et al. Germline SAMD9 and SAMD9L mutations are associated with extensive genetic evolution and diverse hematologic outcomes. JCI Insight. 2018;3(14).

22. Allenspach EJ, Soveg F, Finn LS, So L, Gorman JA, Rosen ABI, et al. Germline SAMD9L truncation variants trigger global translational repression. J Exp Med. 2021;218(5).

23. de Jesus AA, Hou Y, Brooks S, Malle L, Biancotto A, Huang Y, et al. Distinct interferon signatures and cytokine patterns define additional systemic autoinflammatory diseases. J Clin Invest. 2020;130(4):1669–82.

24. Russell AJ, Gray PE, Ziegler JB, Kim YJ, Smith S, Sewell WA, et al. SAMD9L autoinflammatory or ataxia pancytopenia disease mutations activate cell-autonomous translational repression. Proc Natl Acad Sci U S A. 2021;118(34).

25. Thomas ME, 3rd, Abdelhamed S, Hiltenbrand R, Schwartz JR, Sakurada SM, Walsh M, et al. Pediatric MDS and bone marrow failure-associated germline mutations in SAMD9 and SAMD9L impair multiple pathways in primary hematopoietic cells. Leukemia. 2021;35(11):3232–44.

26. Mekhedov SL, Makarova KS, Koonin EV. The complex domain architecture of SAMD9 family proteins, predicted STAND-like NTPases, suggests new links to inflammation and apoptosis. Biol Direct. 2017;12(1):13.

27. Peng S, Meng X, Zhang F, Pathak PK, Chaturvedi J, Coronado J, et al. Structure and function of an effector domain in antiviral factors and tumor suppressors SAMD9 and SAMD9L. Proc Natl Acad Sci U S A. 2022;119(4).

28. Ray S, Hewitt K. Sticky, Adaptable, and Many-sided: SAM protein versatility in normal and pathological hematopoietic states. Bioessays. 2023;45(8):e2300022.

29. Zhou M, Li Y, Hu Q, Bai XC, Huang W, Yan C, et al. Atomic structure of the apoptosome: mechanism of cytochrome c- and dATP-mediated activation of Apaf-1. Genes Dev. 2015;29(22):2349–61.

30. Chen K, Wang D, Kurgan L. Systematic investigation of sequence and structural motifs that recognize ATP. Comput Biol Chem. 2015;56:131–41.

31. Longo LM, Jablonska J, Vyas P, Kanade M, Kolodny R, Ben-Tal N, et al. On the emergence of P-Loop NTPase and Rossmann enzymes from a Beta-Alpha-Beta ancestral fragment. Elife. 2020;9.

32. Demkiv AO, Toledo-Patino S, Medina-Carmona E, Berg A, Pinto GP, Parracino A, et al. Redefining the Limits of Functional Continuity in the Early Evolution of P-Loop NTPases. Mol Biol Evol. 2025;42(4).

33. Krissinel E, Henrick K. Inference of macromolecular assemblies from crystalline state. J Mol Biol. 2007;372(3):774–97.

34. Zhang JT, Liu XY, Li Z, Wei XY, Song XY, Cui N, et al. Structural basis for phage-mediated activation and repression of bacterial DSR2 anti-phage defense system. Nat Commun. 2024;15(1):2797.

35. Yin H, Li X, Wang X, Zhang C, Gao J, Yu G, et al. Insights into the modulation of bacterial NADase activity by phage proteins. Nat Commun. 2024;15(1):2692.

36. Shen Z, Lin Q, Yang XY, Fosuah E, Fu TM. Assembly-mediated activation of the SIR2-HerA supramolecular complex for anti-phage defense. Mol Cell. 2023;83(24):4586–99 e5.

37. Pastor VB, Sahoo SS, Boklan J, Schwabe GC, Saribeyoglu E, Strahm B, et al. Constitutional SAMD9L mutations cause familial myelodysplastic syndrome and transient monosomy 7. Haematologica. 2018;103(3):427–37.

38. Legrand A, Demeure R, Chantharath A, Rey C, Baltenneck J, Gilchrist CLM, et al. Ancient convergence with prokaryote defense and recent adaptations to lentiviruses in primates characterize the ancestral immune factors SAMD9s. bioRxiv. 2025.

39. Leipe DD, Koonin EV, Aravind L. STAND, a class of P-loop NTPases including animal and plant regulators of programmed cell death: multiple, complex domain architectures, unusual phyletic patterns, and evolution by horizontal gene transfer. J Mol Biol. 2004;343(1):1–28.

40. Cheng TC, Hong C, Akey IV, Yuan S, Akey CW. A near atomic structure of the active human apoptosome. Elife. 2016;5.

41. Gao LA, Wilkinson ME, Strecker J, Makarova KS, Macrae RK, Koonin EV, et al. Prokaryotic innate immunity through pattern recognition of conserved viral proteins. Science. 2022;377(6607):eabm4096.

42. Wang Y, Tian Y, Yang X, Yu F, Zheng J. Filamentation activates bacterial Avs5 antiviral protein. Nat Commun. 2025;16(1):2408.

43. Sanders BD, Jackson B, Marmorstein R. Structural basis for sirtuin function: what we know and what we don’t. Biochim Biophys Acta. 2010;1804(8):1604–16.

44. Zhao H, Brautigam CA, Ghirlando R, Schuck P. Overview of current methods in sedimentation velocity and sedimentation equilibrium analytical ultracentrifugation. Curr Protoc Protein Sci. 2013;Chapter 20:Unit20 12.

45. Schuck P. Size-distribution analysis of macromolecules by sedimentation velocity ultracentrifugation and lamm equation modeling. Biophys J. 2000;78(3):1606–19.

46. Brown PH, Schuck P. Macromolecular size-and-shape distributions by sedimentation velocity analytical ultracentrifugation. Biophys J. 2006;90(12):4651–61.

47. Brautigam CA. Calculations and Publication-Quality Illustrations for Analytical Ultracentrifugation Data. Methods Enzymol. 2015;562:109–33.

48. Punjani A, Rubinstein JL, Fleet DJ, Brubaker MA. cryoSPARC: algorithms for rapid unsupervised cryo-EM structure determination. Nat Methods. 2017;14(3):290–6.

49. Tan YZ, Baldwin PR, Davis JH, Williamson JR, Potter CS, Carragher B, et al. Addressing preferred specimen orientation in single-particle cryo-EM through tilting. Nat Methods. 2017;14(8):793–6.

50. Jumper J, Evans R, Pritzel A, Green T, Figurnov M, Ronneberger O, et al. Highly accurate protein structure prediction with AlphaFold. Nature. 2021;596(7873):583–9.

51. Pettersen EF, Goddard TD, Huang CC, Meng EC, Couch GS, Croll TI, et al. UCSF ChimeraX: Structure visualization for researchers, educators, and developers. Protein Sci. 2021;30(1):70–82.

52. Emsley P, Cowtan K. Coot: model-building tools for molecular graphics. Acta Crystallogr D Biol Crystallogr. 2004;60(Pt 12 Pt 1):2126–32.

53. Adams PD, Afonine PV, Bunkoczi G, Chen VB, Davis IW, Echols N, et al. PHENIX: a comprehensive Python-based system for macromolecular structure solution. Acta Crystallogr D Biol Crystallogr. 2010;66(Pt 2):213–21.

54. Williams CJ, Headd JJ, Moriarty NW, Prisant MG, Videau LL, Deis LN, et al. MolProbity: More and better reference data for improved all-atom structure validation. Protein Sci. 2018;27(1):293–315.

55. Schrodinger, LLC. The PyMOL Molecular Graphics System, Version 1.8. 2015.

56. Knight, MJ, Leettola, C, Gingery, M, Li, H & Bowie, JU. A human sterile alpha motif domain polymerizome. Protein Sci. 2011, 20 (10):697–1706.

